# FAIRTraits: an environmentally, methodologically and semantically enriched database of plant traits from Mediterranean populations of 240 species

**DOI:** 10.1101/2025.03.06.641916

**Authors:** Éric Garnier, Léo Delalandre, Jules Segrestin, Karim Barkaoui, Elena Kazakou, Marie-Laure Navas, Denis Vile, Cyrille Violle, Astrid Bellmann, Maud Bernard-Verdier, Marine Birouste, Alain Blanchard, Iris Bumb, Pablo Cruz, Sandrine Debain, Adeline Fayolle, Claire Fortunel, Karl Grigulis, Gérard Laurent, Sandra Lavorel, Francisco Lloret, Ignacio M. Pérez-Ramos, Iván Prieto, Jean Richarte, Catherine Roumet

## Abstract

Trait-based approaches have proven, and continue to offer strong potential to tackle key issues in ecology, including: (i) understanding the functioning of organisms and how it relates to the environment, (ii) identifying the rules governing the assembly of communities and the coexistence of species, (iii) understanding how the functioning of organisms scales up to that of ecosystems and controls some of the services they deliver to humans. We present FAIRTraits, a data set of plant traits assembled from a set of studies designed to address these issues in the pedo-climatic context of the Northern Mediterranean Basin, considered both a biodiversity and a climatic hotspot. The FAIR (Findable, Accessible, Interoperable and Reusable) guiding principles were followed to ensure maximum visibility and reusability of these data.

FAIRTraits compiles and standardizes trait data collected over the 1997-2023 period on six sites by the same research group, ensuring that a consistent methodology was followed. These data were obtained on individuals from 1,955 populations of 240 species belonging to 155 genera and 48 families (172 herbaceous and 39 woody species). It contains 189,452 records for 183 quantitative traits from the different plant organs (leaves, stems, roots, and reproductive parts), which have been grouped into 10 categories: allocation ratio, architecture, chemistry, dynamics, mechanics, microbial associations, morphology, phenology, physiology and plant size. Trait values are given at the level of individual measurements. Species-level values of height and phenology taken from a Mediterranean flora are also given. Species are characterized by plant family, life cycle, Raunkiaer lifeform, photosynthetic pathway and by an original successional stage indicator value. As trait values strongly depend on environmental conditions, we also provide information on the climatic conditions and soil properties of the sites, as well as on disturbance regimes of the plots in which sampled individuals were collected.

The following steps were taken to ensure the FAIRness of the data set. *Findable*: metadata are described using the Ecological Metadata Language, and are deposited both on the InDoRES (CNRS/MNHN) metadata catalogue and on the GBIF data portal (see below); *Accessible*: FAIRTraits is available on the InDoRES repository, with a subset available on the GBIF data portal (see below); *Interoperable*: recognized taxonomical and terminological resources to qualify taxa and attributes have been used whenever possible, and fully described sampling protocols and measurement methods are given both for traits and environmental data. A subset of the data could be mapped onto the Darwin Core biodiversity standard, making it possible to display part of the data set on the GBIF portal, more traditionally used for taxonomic occurrence data. *Reusable*: (meta)data are thoroughly described using domain-relevant community standards, and the full data set is released under the CC-BY 4.0 license.

We believe that these multiple efforts, spanning from the very content of the data set to its careful formatting, will make of FAIRTraits a highly valuable resource for trait-based research, both in terms of data analysis and reusability.

## Class I. Data Set Descriptors

### A. Data set identity

FAIRTraits: an environmentally, methodologically and semantically enriched database of plant traits from 1,955 Mediterranean populations of 240 species belonging to 155 genera and 48 families.

### B. Data set identification code

The FAIRTraits database is composed of nine csv files listed in Table 1. Metadata and data have been split into several files corresponding to different aspects of the database.

**Table 1.** Main elements of the FAIRTraits database, with their type (data or metadata) and a short description of their contents and state. See further tables below for details on each component.

| Element | Type | Destination | Contents |
| --- | --- | --- | --- |
| FAIRTraits_Authors.csv | Metadata | InDoRES, GBIF | information on contributors to the database |
| FAIRTraits_Projects.csv | Metadata | InDoRES, GBIF | information on projects during which data were collected |
| FAIRTraits_Taxon.csv | Data | GBIF (Darwin Core archive) | information on taxa found in the database with selected characteristics |
| FAIRTraits_TraitValues.csv | Data | GBIF (Darwin Core archive) | the core file with trait values and relevant information |
| FAIRTraits_MeteoSitesMonth.csv | Data | InDoRES | data on monthly averages of meteorological variables from the meteorological stations closest to the six sites |
| FAIRTraits_MeteoSitesYear.csv | Data | InDoRES | data on yearly averages of meteorological variables from the meteorological stations closest to the six sites |
| FAIRTraits_EnvironmentalPlots.csv | Data | InDoRES | information on georeferencing, treatments, type of soils and soil properties of plots in which environmental data were collected |
| FAIRTraits_DescriptionEnvVariables.csv | Metadata | InDoRES | a description of variables, methods and items found in files FAIRTraits_MeteoSitesMonth.csv, FAIRTraits_MeteoSitesYear.csv;<br>FAIRTraits_EnvironmentalPlots.csv, |
| FAIRTraits_PublishedPapers.csv | Metadata | InDoRES | a list of scientific articles in which subsets of the database have been used |

### C. Data set description

#### 1. Originators

Éric Garnier, Marie-Laure Navas, Elena Kazakou, Cyrille Violle and Catherine Roumet

#### 2. Abstract

FAIRTraits compiles and standardizes trait data collected by the same research group, to address a range of topics including the study of: (1) structure-function relationships in plants, (2) trait-environment relationships, (3) scaling from plant traits to ecosystem properties, (4) trait-demography relationships and (5) phylogenetic signal in plant traits. Traits were collected from six Mediterranean sites over the period 1997-2023 from 240 species belonging to 155 genera and 48 families. It contains 189,452 records for 188 quantitative traits, which can be broadly categorized into: (1) morpho-anatomical leaf (area, thickness, dry matter content, specific leaf area, etc.) and root (diameter, specific root length, dry matter content, etc.) traits, (2) chemical leaf, stem and root traits (carbon, nitrogen, phosphorus contents, carbon isotopic fraction, etc.), (3) whole plant and reproductive traits (vegetative and reproductive heights, timing of reproductive events, seed mass and number), (4) physiological traits (photosynthetic rate, stomatal conductance, root respiration, etc.) and (5) mass loss rates during decomposition of leaves, stems and roots. Trait values are given at the level of individual replicate measurements, without any aggregation. Species-level values of height and phenology taken from a Mediterranean flora are also given. Species are characterized by plant family, life cycle, Raunkiær lifeform, photosynthetic pathway and by an original successional stage indicator value. As trait values strongly depend on environmental conditions, we also provide information on the climatic conditions and soil properties of the sites, as well as on disturbance regimes of the plots in which sampled individuals were collected.

A number of steps, described in the different sections below, were taken to ensure that the database follows the FAIR guiding principles (Findable, Accessible, Interoperable and Reusable) for scientific data management and stewarship (Wilkinson et al. 2016).

### D. Key words/phrases

biodiversity standards; ecosystem properties; environmental conditions; Mediterranean Biogeographic Region; leaf, stem and root traits; plant functional traits; gas exchange; litter mass loss; phenology; reproductive traits; terminological resources; trait-based ecology

## Class II. Research origin descriptors

### A. Overall project description

#### 1. Identity

FAIRTraits compiles and standardizes data collected on six sites located at the north-western edge of the Mediterranean Biogeographic Region, in southern France (five sites) and north-eastern Spain (one site), during the course of 12 research projects (see Table 2 in section Class II.A.5.) conducted within the same research group at CEFE.

**Table 2.** Research projects during which data were collected, with their principal investigator, the period and source of funding. Details on the specific objectives of each project can be found in the FAIRTraits_Projects.csv file, one of the FAIRTraits basic elements (see Table 1)

| Project acronym | Full name of project | Principal investigator | Period of study | Source of funding |
| --- | --- | --- | --- | --- |
| MELODY | Mediterranean Landscapes in a Changing World: Coupling Dynamics and Functional Analyses | Serge Rambal | 1997-2003 | ICSU – IGBP – Core Research Project |
| DynEcoMed | Dynamique des Ecosystèmes Méditerranéens dans un Monde Changeant | Sandra Lavorel | 2001-2006 | CNRS – Laboratoire Européen Associé CEFE-CREAF |
| INDIGO | Une Nouvelle Perspective sur les Indicateurs de Diversité Végétale : Application à l'Etude des Conséquences de la Déprise Agricole et des Valeurs d'Usage des Prairies | Eric Garnier | 2002-2006 | MATE – Programme « Action Publique, Agriculture et Biodiversité » |
| VISTA | Vulnerability of Ecosystem Services to Land Use Change in Traditional Agricultural Landscapes | Sandra Lavorel | 2002-2006 | European Union – Contract n°EVK2-2001-000356 |
| GEOTRAITS | Impacts des Traits Fonctionnels des Espèces Végétales sur les Cycles Biogéochimiques | Eric Garnier | 2004-2007 | ACI – FNS Ecosphère Continentale (PNBC) |
| DivHerbe | Structure, Diversité et Fonctionnement : des Clefs Multi-Echelles pour la Gestion des Prairies Permanentes | Eric Garnier | 2006-2009 | INRA – AIP EcoGer |
| RESPIRS | The Root Economics Spectrum and InfraRed Spectroscopy | Catherine Roumet | 2010-2012 | FRB – Programme « Petits et percutants » |
| O2LA | Organismes et Organisations Localement Adaptés | Marie-Laure Navas | 2010-2015 | ANR – Contract n°09-STRA-09 |
| CASCADE | Interactions Trophiques et Fonctionnement des Ecosystèmes Terrestres : une Approche Fonctionnelle et Chimique | Eléna Kazakou | 2013-2014 | CNRS – INSU – InEE – INC – Initiative structurante EC2CO |
| PhenBiom | Phenology and Biomechanics of Plants in the Mediterranean | Eric Garnier | 2013-2018 | CNRS – Core support funding |
| COMODO | Connecter les Mondes Verts et Bruns : Ecologie Trophique des Détritivores et des Phytophages en Prairie Permanente | Sylvain Coq | 2015-2017 | CNRS – INSU – Initiative structurante EC2CO |
| FageAnnuals | Functional Ecology of Annual Plant Species from a Mediterranean Rangeland | Cyrille Violle | 2021-2023 | European Union - ERCStG-2014-639706 |

#### 2. Originators

Éric Garnier conceived and coordinated the project. Éric Garnier, Marie-Laure Navas, Catherine Roumet, Elena Kazakou and Cyrille Violle elaborated the various research projects leading to the collection of data. All co-authors provided data which were first assembled in a preliminary data set by Jules Segrestin, who also conducted the outlier detection on the finalized database. Éric Garnier finalized the compilation of data in raw data files, Léo Delalandre elaborated the scripts to format the database in its final form, and Karim Barkaoui conducted the analyses describing the content of the database and elaborated the corresponding illustrations.

#### 3. Period of study

The data were collected between 1997 to 2023, during different discrete measurement campaigns at each site.

#### 4. Objectives

To provide a database of plant traits at the individual replicate level, for a broad range of traits collected on individuals from 1,955 Mediterranean populations of 240 species. Methodological details (sampling protocols and measurement methods) are provided for each record, and the interpretation of trait values will be enhanced by the availability of information on the climatic conditions of the sites, soil properties and on disturbance regimes of the plots in which the sampled individuals were collected. The FAIR (Findable, Accessible, Interoperable and Reusable) guiding principles were followed to ensure maximum visibility and reusability of these data.

#### 5. Abstract

There is a growing consensus that a trait-based approach has a strong potential to tackle key issues in ecology, including: (i) understanding the functioning of organisms and how it relates to the environment, (ii) identifying the rules governing the assembly of communities, (iii) understanding how the functioning of organisms scales up to that of ecosystems and controls some of the services they deliver to humans. FAIRTraits is a database of plant traits assembled from various studies designed to address these issues in the pedo-climatic context of the north western Mediterranean Basin, considered both a biodiversity and a climatic hotspot. A short summary of the rationale underlying the projects during which data were collected is given below (see also Table 2 in section Class II.A.5.):

***MELODY*** was a unifying program focusing on biodiversity dynamics at the regional scale of the French Mediterranean area. In this context, FAIRTraits data pertain to post-cultural plant successions.

***DynEcoMed*** was aimed at understanding and predicting the dynamics of vegetation structure and functioning in Mediterranean ecosystems undergoing land use and climate changes.

***INDIGO*** aimed to (i) analyze the consequences of the decrease in land use by agriculture in some areas on biological diversity; (ii) relate these changes in diversity to the functioning of ecosystems and services delivered when relevant; and (iii) propose a new methodology to assess the effects of changes in agricultural practices on vegetation.

***VISTA*** aimed to produce an integrated assessment of the vulnerability of European traditional agro-pastoral landscapes to land use and climate change to assist land managers and regional policy makers towards sustainable development.

***GEOTRAITS*** objectives were to analyze the impacts of differences in plant functional characteristics as assessed through the values of their traits, on biogeochemical cycles.

***DivHerbe*** objectives were to understand the mechanisms that control species diversity and structural heterogeneity in plant communities, and propose simple indicators aimed at managing grassland diversity according to combined environmental and production objectives.

***RESPIRS*** objectives were to test on a large number of temperate, Mediterranean and tropical species: i) the existence of a root economics spectrum consisting of key chemical and structural traits enabling the prediction of root respiration and decomposition, and ii) the potential of near infrared spectroscopy to characterize root diversity.

***O2LA*** was an interdisciplinary research project aiming to characterize the added value of diversification (of biological resources, management practices, knowledge transfers) in French livestock production systems.

***CASCADE*** was designed to understand the response of the soil-plant system to herbivory and identify the consequences of plant responses on the quality of the resources for herbivores on the one hand, and on decomposers through the litter produced on the other hand.

***PhenBiom*** was a research project aiming at integrating plant reproductive phenology and leaf mechanical defense into the phenotypic space of Mediterranean plants.

***COMODO*** objectives were to describe, understand and predict the structure of trophic networks through the identification of relevant trophic interaction traits. Two coupled ecological networks were studied: plants-herbivores and litter-decomposers.

***FageAnnuals*** was aimed at testing whether species differing in their life history display contrasted trait-environment relationships using the case of annual and perennial species, which often co-occur in communities, but whose life cycles fundamentally differ.

#### 6. Sources of funding

Table 2 below lists the original projects during which the data were collected.

### B. Specific subproject description

#### 1. Site description

Data were collected in six sites located at the north-western edge of the Mediterranean Biogeographic Region (Figure 1): five in southern France (Cazarils, Hautes Garrigues du Montpelliérais, La Fage, Les Agros and Camp Redon), and one in north-eastern Spain (Garraf).

**Figure 1.**
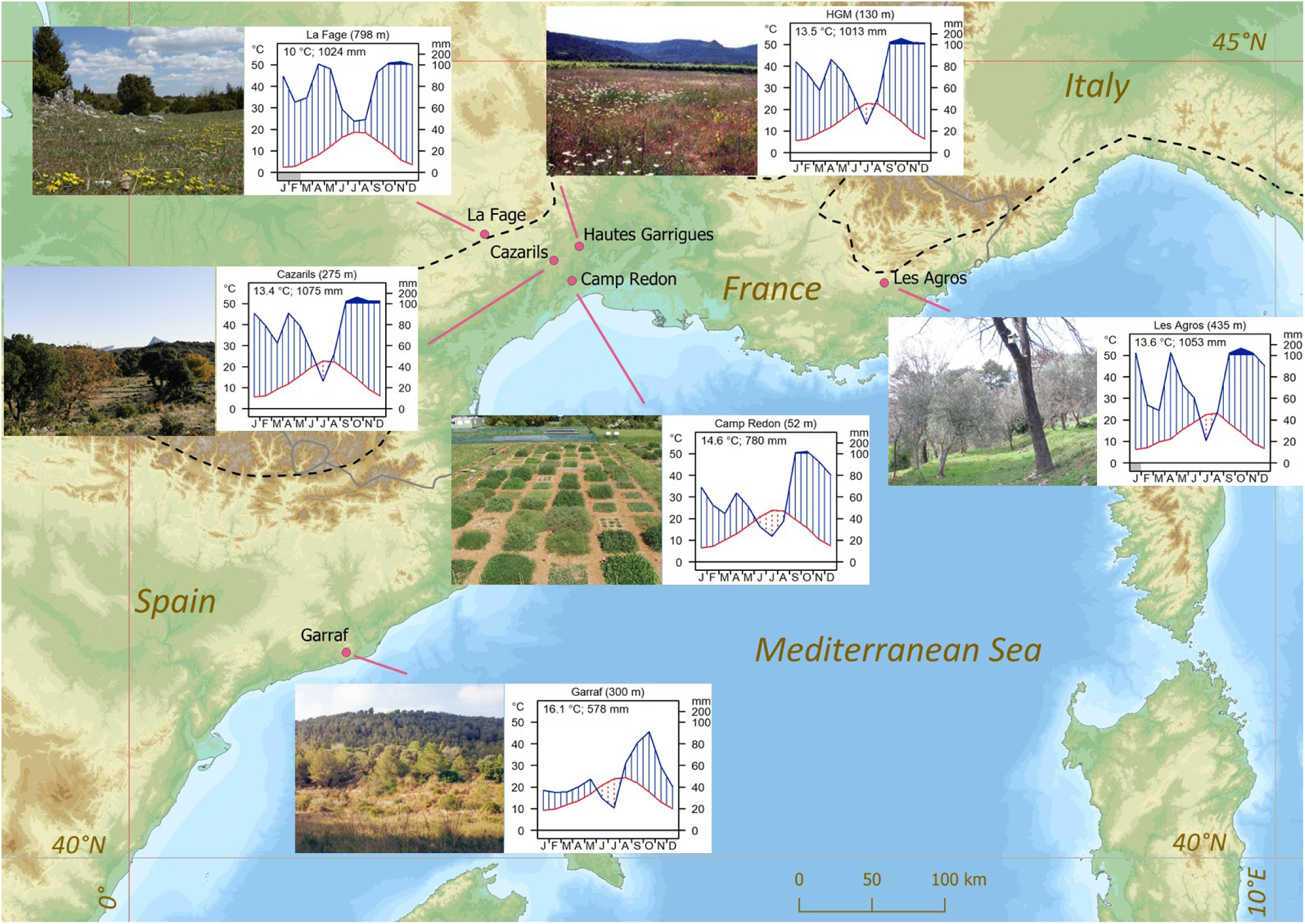
Map showing the location of the six sites where data have been collected. Views of representative areas of the six sites with the corresponding climate diagrams (cf. Walter et al. 1975; blue line: rainfall, red line: temperature) are added. On these diagrams, dry periods correspond to the area with red dots (monthly rainfall <2 × average monthly temperature: cf. Walter et al., 1975). At the bottom of each plot, a grey bar represents months with mean minimum temperatures below 0°C. Table 3 for further details. Colors on the background map correspond to altitudinal variations from low (green) to high (brown) altitudes. The black dashed line shows the northern limit of the Mediterranean Biogeographic Region (cf. Tison et al. 2014).

**Table 3.**
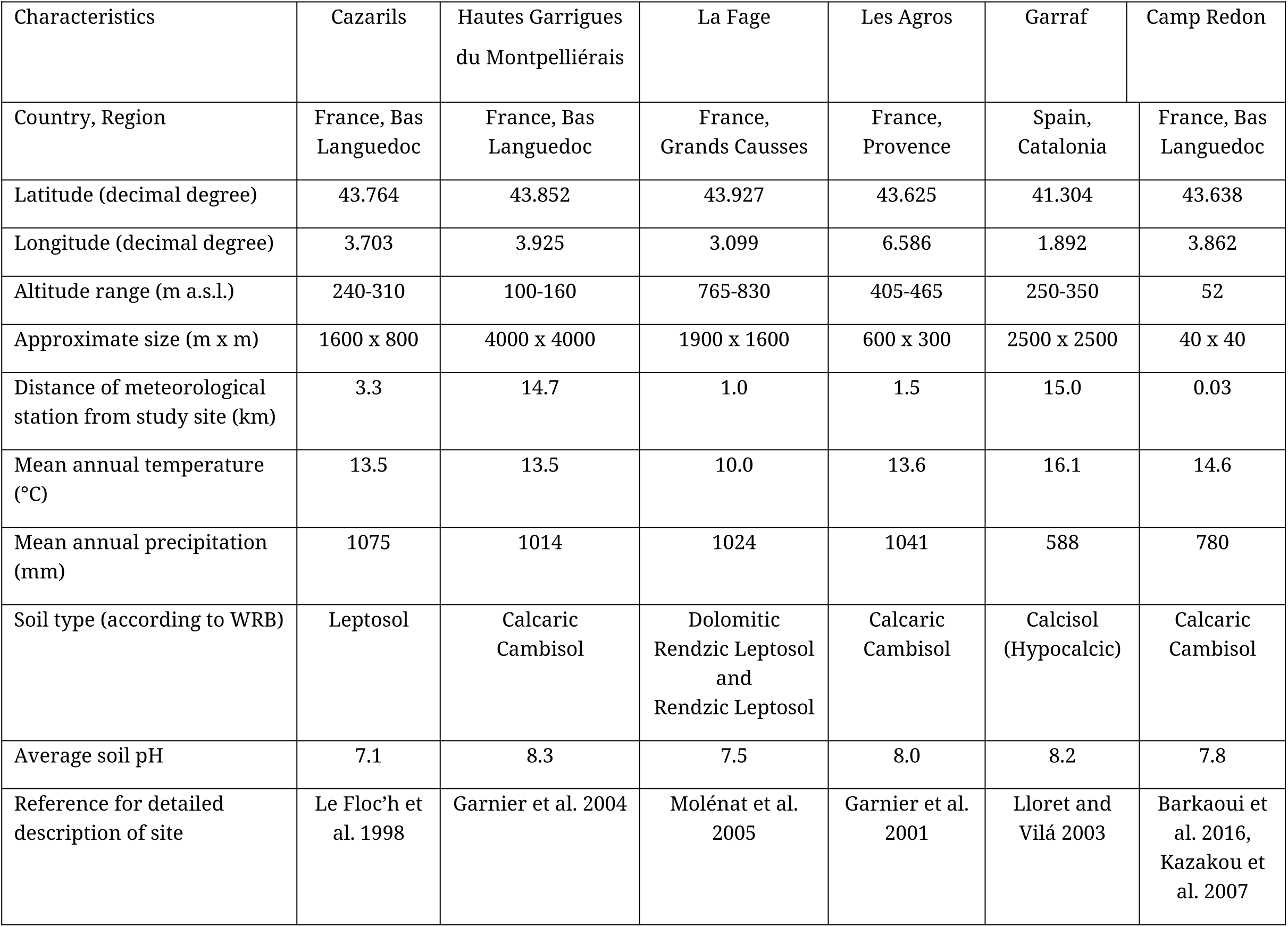
Selected characteristics of the six study sites where data have been collected. Temperature and precipitation data were taken from the meteorological station closest to each site. Soil type follows the WRB classification (World Reference Base for Soil Resources 2022). More details can be found in the FAIRTraits_EnvironmentalPlots.csv, FAIRTraits_MeteoSitesMonth.csv and FAIRTraits_MeteoSitesYear.csv files.

All sites are located *in natura* except Camp Redon, which is an experimental garden. Selected characteristics of these sites are given in Table 3, and views of representative areas are shown in Figure 1. At all sites, the climate is classified as Mediterranean, with mild winters and (relatively) dry summers: it spans from “dry sub-humid” in Garraf, to “humid” in Cazarils, Hautes Garrigues, Les Agros, and Camp Redon, and while still classified as Mediterranean, the climate in La Fage is at the limit of the temperate zone (cf. Figure 1; see Daget 1977 for bioclimate categories). Sites have undergone a strong and long-lasting human imprint (see short description below). The vegetation is generally engaged in highly dynamic processes resulting from changes in land use (e.g. abandonment of cultivation, grazing, fertilization), resulting in a complex mosaic of herbaceous and woody components (Figure 1). In the six sites, the bedrock is limestone and soils show neutral to slightly basic pH.

A short description of each site is given below.

***Cazarils***, a part of the “Domaine de Roussières”, consists of a large clearing in an otherwise mostly wooded area, with *Quercus pubescens* and *Q. ilex* as dominant tree species. This clearing, which was cultivated until the mid-20^th^ century, is surrounded by degraded Mediterranean shrublands (“garrigue”) that were regularly cut for fuel wood collection. Over the past 80 years or so, recreational hunting, selective clearcutting and burning, and extensive grazing by sheep and goats have been the main disturbances at this site. Former and current land uses have resulted in a complex spatial organization of the vegetation with plant communities composed of species from different growth and life forms (Le Floc’h et al. 1998, Navas et al. 2010).

***Hautes Garrigues du Montpelliérais*** consists of a collection of twelve old-fields located in a matrix of cultivated vineyards, other less represented crops and woodlands. These fields were all previously vineyards, which, following removal of the vines, were abandoned 2 to 42 years prior to 2000. The vegetation shifts from predominantly annual species in recently abandoned fields (less than 5 years) to herbaceous perennials in fields of intermediate age (6-15 years), and to a mixture of herbaceous and woody perennials in fields older than 25 years (see Garnier et al. 2004 for details).

***La Fage*** is an experimental farm managed by the French National Research Institute for Agriculture, Food and the Environment (INRAE). The area was probably cultivated until the 19^th^ century, at least partially. It was then extensively grazed until 1978, both by local and transhumant sheep herds, allowing woody species to recolonize a large part of the area. Since 1978, it has been subdivided into paddocks either grazed by sheep at a low stocking rate, or fertilized and grazed by sheep at a high stocking rate, or fenced to prevent grazing. The vegetation consists primarily in rangelands dominated by perennial grasses and forbs, with a small woody component covering approximately 20– 25% of the area (see Molénat et al. 2005 for details).

***Les Agros*** consists of a series of south facing terraces located on the southern edge of a large limestone plateau, which were formerly cultivated mainly with olive trees until the second half of the 19^th^ century. Until approximately 40 years ago, the terraces were occasionally grazed by sheep. Since then, they have been regularly cleared. The vegetation is mostly herbaceous with numerous shrubs and olive tree resprouts, surrounded by a mixed *Pinus halepensis* – *Quercus ilex* woodland (see Garnier et al. 2001 for details).

***Garraf*** (Garraf Natural Park) is a karstic massif where the vegetation is a mosaic of shrublands (including “garrigue”) and *Pinus halepensis* open forests growing on rocky soils and on terraces that were established over marls and cultivated in the past centuries. Agricultural abandonment started at the end of the 19^th^ century and was completed around the mid-20^th^ century. Several wildfires have occurred since then, particularly in 1982 and 1994, affecting most of the area. Shrubs (macrophanerophytes), scrubs (nanophanerophytes) and herbaceous species are important components of the community (see Lloret and Vilá 2003 for details).

***Camp Redon*** is a 2.5 ha experimental garden attached to the CEFE research unit, located in the outskirts of the Montpellier city. Data for FAIRTraits are derived from two experiments conducted on small areas of this garden. Plants were cultivated in plots previously weeded and in which the 5cm upper layer of soil had been removed. The first experiment (“PDM”) was conducted in 2004 and 2005 on 18 species from the old-field succession of the Hautes Garrigues du Montpelliérais site described above, grown under high and low nitrogen supply (Kazakou et al. 2007 for details). The second one (“O2LA”) was conducted in 2012 and 2013 on a selection of perennial species from the La Fage Mediterranean rangeland described above, grown at two levels of water supply (Barkaoui et al. 2016; Fort et al. 2017 for details).

#### 2. Experimental or sampling design

##### a. Design characteristics

***Spatial design:*** in the case of the five *in natura* sites, delimited topographic zones of various areas named “plots” were recognized, which are the spatial units in which individuals were selected for trait measurement (114 plots across all sites). Each plot was assigned to a single modality of a specific treatment (taken in a broad sense), which differ in each site. In the case of Camp Redon, there were 144 plots (1.20 x 1.20 m) overall (72 at each level of nitrogen supply) in the PDM experiment, and 76 plots (1 x 1.20 m) overall (64 and 12 in the “ambient” and “low” water supplies, respectively) in the O2LA experiment. Each trait record in the database is attached to one and only one of these plots.

***Species selection for trait measurements:*** measurements were conducted on the most abundant species from each site. When the objective was to test the effects of community functional structure on ecosystem properties, we sampled species that collectively made up at least 80% of vegetation cover and/or biomass (see Garnier et al. 2004, Pakeman & Quested 2007). In the case of the two experiments conducted in Camp Redon, species were selected on the basis of their frequency in the different stages of the chrono sequence studied in Hautes Garrigues du Montpelliérais (PDM), and of their abundance in the low stocking rate, unfertilized treatment in La Fage (O2LA).

Traits were measured on 172 herbaceous (71 therophytes, 87 hemicryptophytes and 14 geophytes) and 69 woody (35 chamaephytes, 11 deciduous and 18 evergreen phanerophytes, and 5 lianas) species. Asteraceae (36 species), Poaceae (34 species), Fabaceae (29 species), Lamiaceae (16 species) and Rosaceae (14 species) are the five most represented botanical families in the database. With one exception (Rosaceae), this corresponds to the representation of families in BROT 2.0, a database of plant functional traits across the Mediterranean Basin (Tavşanoğlu and Pausas 2018). The five most represented species are *Bromopsis erecta* (Poaceae, 11,541 records), *Potentilla neumanniana* (Rosaceae, 6748 records), *Sanguisorba minor* (Rosaceae, 4204 records), *Teucrium chamaedrys* (Lamiaceae, 4120 records), and *Arenaria serpyllifolia* (Caryophyllaceae, 3365 records). Figure 2 gives the number of species and populations measured in each site, and the number of populations measured for the most represented species in the database.

**Figure 2.**
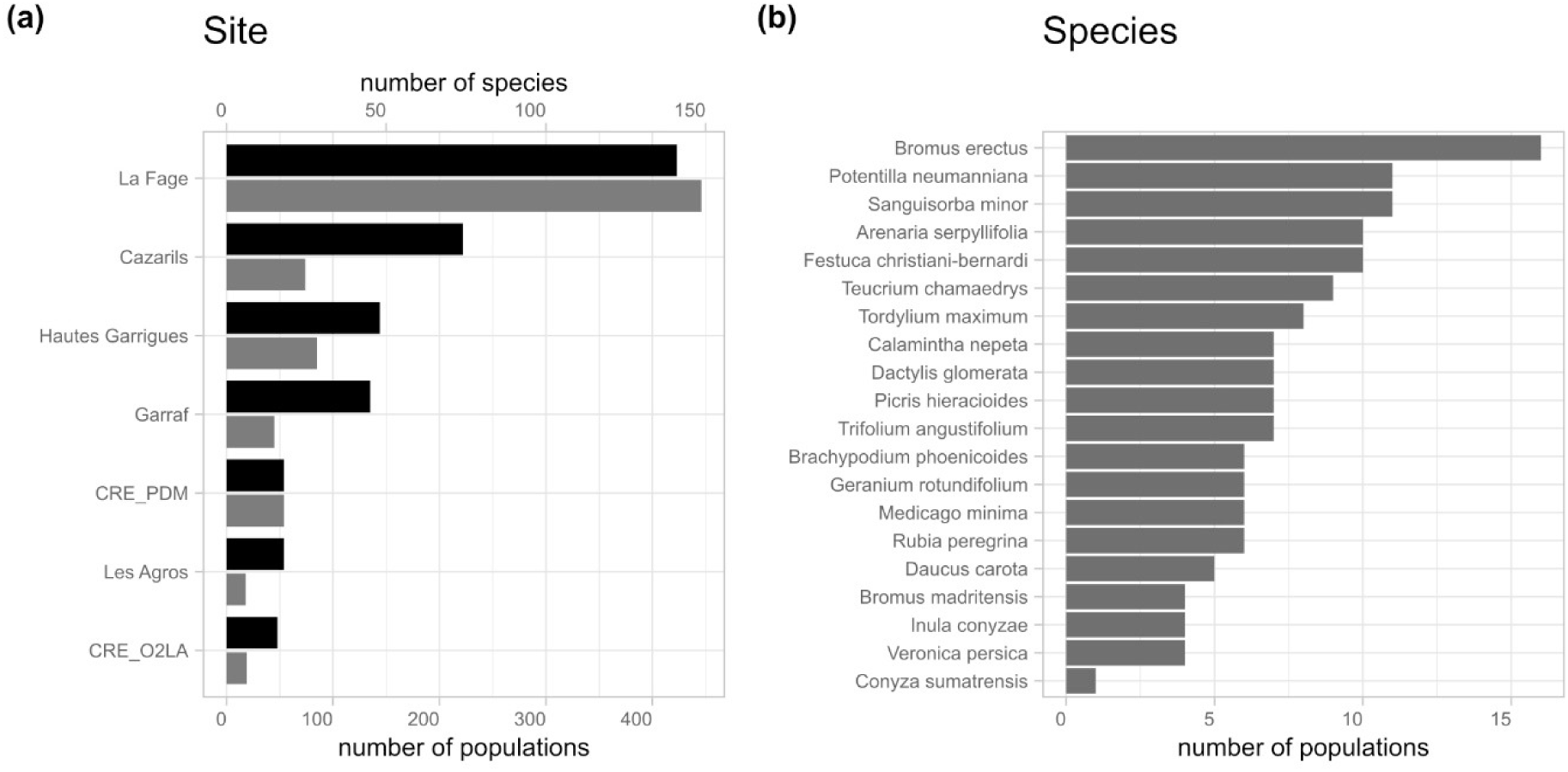
(a) cumulated number of species (black bars, upper x-scale) and populations (grey bars lower x-scale), grouped by site, and (b) cumulated number of populations for the 20 most represented species in the FAIRTraits database.

***Environmental factors:*** environmental data are provided at the site and/or at the plot level. At the site level, 30-year average values of a number of monthly and yearly climatic variables were compiled from meteorological stations closest to the sites. Within each site, the local environment was more precisely characterized in a subset of 59 plots, selected to represent either a specific treatment or a particular topographic zone: disturbance type, frequency and intensity, soil type (cf. Table 3 in section Class II.B.1.) and 11 quantitative soil variables were quantified in these plots. Each plot in which traits were assessed was then associated with one of these “environmental plots”.

##### b. Data collection period

Across the whole database, data collection took place between 1997 and 2023. The main sampling periods differ among sites, in relation with the sites selected for the various projects listed in Table 2 (section Class II.A.6.): 1997-2001 in Cazarils with some additional sampling in 2017 and 2023; 1998-1999 in Les Agros; 2000-2004 in Hautes Garrigues du Montpelliérais; 2001 in Garraf; 2006-2022 in La Fage; 2004-2006 (PDM) and 2012-2013 (O2LA) in Camp Redon. Samples of plant vegetative organs were mostly collected between March and July, corresponding to the main growing season under the Mediterranean climate prevailing at the different sites. Seed production and seed traits were assessed when seeds were ripe, while plant litter was collected at peak litter production, in the summer and early autumn. The exact date at which measurements were taken is provided for 97% of the data in the FAIRTraits_TraitValues.csv file.

#### 3. Research methods

For each record in the FAIRTraits_TraitValues.csv file database, a short description of the sampling protocol (field “samplingProtocol”) and of the method used to determine the trait value with relevant references (field “measurementMethod”) are given (see Table 4 in section Class IV.B.1. for the structure of this file). This was also done for the environmental data in the three corresponding files (see Table 7 in section Class IV.B.4.). Given the large number of traits measured (179 overall: section Class IV.B.2.), we give only a short overview of methods in this section, and direct the reader to the relevant data files for further details.

**Table 4.** Structure of the FAIRTraits_Taxon.csv file. The Darwin Core terms and classes are given for each variable, when available. Units, literature sources and methods are given in the csv file.

| Name of variable | Variable definition | Darwin Core term | Darwin Core class |
| --- | --- | --- | --- |
| **SpeciesName** | Name of species taken from a local flora (latin binomial without authority) | acceptedNameUsage | Taxon |
| **taxonRank** | The taxonomic rank of the most specific name in the Darwin Core field “scientificName” | taxonRank | Taxon |
| **CodeSp** | Identification code of the species (4 first letters of genus name and 4 first letters of species name) | taxonID | Taxon |
| **Kingdom** | The full scientific name of the kingdom in which the taxon is classified | kingdom | Taxon |
| **Family** | The botanical family of the species | family | Taxon |
| **LifeCycle1** | Primary life cycle of species (annual, monocarpic perennial, perennial) | LifeCycleUnstructured | Plinian Core Simple Extension |
| **LifeCycle2** | Secondary life cycle of species (annual, bisannual, monocarpic perennial, perennial), if any | NA | NA |
| **LifeForm1** | Primary life form of species *sensu* Raunkiaer (1934) | LifeFormUnstructured | Plinian Core Simple Extension |
| **LifeForm2** | Secondary life form of species *sensu* Raunkiaer (1934), if any | NA | NA |
| **SuccessionalStage** | Successional Stage Indicator Value | HabitatUnstructured | Plinian Core Simple Extension |
| **PlantHeight** | Minimum plant height - Maximum plant height (cm) | NaturalHistoryUnstructured | Plinian Core Simple Extension |
| **FloweringStart** | Beginning of flowering period (month) | StartTimeInterval | Plinian Core Simple Extension |
| **FloweringEnd** | End of flowering period (month) | EndTimeInterval | Plinian Core Simple Extension |
| **PhotosyntheticPathway** | Primary mode of atmospheric carbon dioxide fixation (C<sub>3</sub>, C<sub>4</sub>, CAM) | TrophicStrategy | Plinian Core Simple Extension |
| **Chorology** | Geographic distribution of the species | DistributionUnstructured | Plinian Core Simple Extension |
| **scientificName** | Scientific name of the species with authority | scientificName | Taxon |
| **nameAccordingTo** | Identifiant number of the species in TaxRefv16 | nameAccordingTo | Taxon |
| **scientificNameID** | Uniform Resource Identifier of the species in TaxRefv16 | scientificNameID | Taxon |

##### a. Field/laboratory

***Traits:*** for above-ground vegetative and reproductive traits, sampling and measurement methods generally follow Cornelissen et al. (2003), a handbook completed during the period of the first research projects which led to the collection of data synthesized in FAIRTraits. For below-ground traits, we mostly followed the “starting guide to root ecology” (Freschet et al. 2021), which assembles methods developed and improved over the last two decades or so. Traits were mostly determined on healthy individuals of the targeted species found in full light conditions. Some traits were measured non-destructively on-site on intact, undisturbed individuals (plant height, lateral spread, reproductive phenology, photosynthetic rate, etc.), while for traits requiring further processing in the laboratory (leaf area and mass, root length and mass, seed shape and mass, leaf and root nutrient content, etc.), as far as possible, material free from herbivore or pathogen damage was collected. The number of samples (individuals or parts thereof) on which traits were determined varied according to the trait (cf. Table 3 in Cornelissen et al. 2003 for above-ground traits). The exact number of replicates used for each trait can be retrieved from the FAIRTraits files, in which the finest level of replication is conserved.

***Environmental data:*** at the site level, climate was described using data from the meteorological station closest to the site (i.e. one value *per site* for each variable retained). In the 59 environmental plots described above (section Class II.B.2.a.), disturbance frequency and intensity were assessed following Garnier et al. (2007), and soil samples from different layers (specified in the FAIRTraits_EnvironmentalPlots.csv) were collected. These samples were sent to the INRAE “Laboratoire d’Analyses des Sols d’Arras” (https://las.hautsdefrance.hub.inrae.fr/) for analyses. A short summary of methods is given in the FAIRTraits_EnvironmentalPlots.csv file for each variable, and a full description can be found in the “catalogue analytique” of the INRAE laboratory available at: https://las.hautsdefrance.hub.inrae.fr/prestations/catalogue-analytique.

##### b. Instrumentation

A whole range of tools has been used to determine the 188 traits of the database: human eyes (phenological records), rulers (plant height, lateral spread, etc.), precision balances (masses), flatbed scanners coupled with image analysis systems (leaf area, root length and diameter, etc.), portable gas exchange systems (photosynthesis, root respiration), elemental (carbon, nitrogen content) and colorimetric analyzers (phosphorus content), mass spectrometers (element isotopic fractions), etc.. In addition, some trait values were derived from calculations based on raw data (rooting depth, litter mass remaining standardized after a given period, etc.). The brand and model of the devices used as well as the calculations procedures when relevant, are given for each record in the “measurementMethod” field of the database (see Table 4 in section Class IV.B.1.).

##### c. Taxonomy and systematics

See section Class IV.B.3. below for taxonomy and additional features retained to describe taxa.

##### d. Permit history

***Cazarils:*** a formal permit was issued by the Hérault Departmental Council, the administrative entity that owns and manages the “Domaine de Roussières”, to conduct field work and collect samples.

***Hautes Garrigues du Montpelliérais:*** the owners of the land where the twelve old-fields were located were contacted and gave verbal permission to conduct field work and collect samples on their properties until the INDIGO and VISTA projects came to an end.

***Les Agros:*** the owner gave a verbal permission to conduct field work and collect samples on this private property. He participated in the collection of leaf life span data at this site, and was offered co-authorship on the related publication.

***Garraf:*** a formal permit was issued by the administration of the Garraf National Park to CREAF to conduct field work and collect samples.

***La Fage***: the “Génétique Animale” department of INRAE, who rules the experimental farm, has given a long-term agreement to CEFE researchers to conduct research in this site. The staff of the farm is regularly associated as co-authors to publications resulting from the work conducted there.

***Camp Redon:*** as an entity attached to the CEFE laboratory, permission to use specific areas of the experimental garden is discussed with the technical staff in charge of its management when required.

##### e. Legal/organizational requirements

all intermediate and final reports of projects listed in Table 2 (section Class II.A.6.) were delivered as required.

#### 4. Project personnel

Co-authors of this data paper, laboratory technicians and undergraduate students from the CEFE and AGIR research units.

## Class III. Data set status and accessibility

### A. Status

1. **Latest update:** 20 December 2024
2. **Latest archive date:** data not archived as yet
3. **Metadata status:** last update on 20 December 2024
4. **Data verification:** see section Class V.B.

### B. Accessibility

#### 1. Storage location and medium

The different files constituting the database are available as Supporting Information with this *Ecology* data paper. Upon acceptance, metadata will be archived in the InDoRES metadata catalog (https://cat.indores.fr/geonetwork/srv/eng/catalog.search#/home) and will be available as an EML file on the GBIF portal; data will be archived in the following repositories: InDoRES (https://www.indores.fr/) and GBIF (https://www.gbif.org/fr/) for data formatted using the Darwin Core data standard (Table 1 in section Class I.B. for details). The associated R code will be available on GitHub (Table 8 in section Class V.C. for details).

#### 2. Contact persons

**a.** *Eric Garnier:*

CEFE, Univ Montpellier, CNRS, EPHE, IRD, Montpellier, France

Phone: (+) 33 4 67 61 32 42

**b.** *Cyrille Violle:*

CEFE, Univ Montpellier, CNRS, EPHE, IRD, Montpellier, France

Phone: (+) 33 4 67 61 33 42

**c.** *Karim Barkaoui:*

CIRAD, UMR AMAP/ABSys, Montpellier, France

Phone: (+) 33 6 75 59 82 40

### 3. Copyright restrictions

The database is licensed under the CC-By Attribution 4.0 International. As such, it can be used for non-commercial purposes.

### 4. Proprietary restrictions

We request that users of the FAIRTraits database or portion thereof to cite this data paper. In addition, when relevant, users may want to cite the original study in which the data have been collected and presented in the first place. A list of papers with associated projects (Table 2 in section Class II.A.6.) is given below (section Class V.E.) and is available in the FAIRTraits_PublishedPapers.csv file.

### 5. Costs

There are no costs associated with using the FAIRTraits database.

## Class IV. Data structural descriptors

### A. Data set files

#### 1. Identity

Following the approach presented by Schneider et al. (2019), data have been divided into five files (see Table 1 in section Class I.B.). A core file (FAIRTraits_TraitValues.csv) containing trait values and contextual information, with four extension files giving detailed information on taxa (FAIRTraits_Taxon.csv), environmental conditions at the plot level (FAIRTraits_EnvironmentalPlots.csv) and climatic variables at the site level (FAIRTraits_MeteoSitesMonth.csv for average monthly data and FAIRTraits_MeteoSitesYear.csv for average yearly data). Additional “metadata files” (listed in Table 1, section Class I.B.) provide detailed descriptions of the fields that make up the database.

#### 2. Size of data files

***FAIRTraits_TraitValues.csv*:** 245,251 rows (excluding the header); 38 columns; 198 Mo. In this core file, the data are in long format where each row represents values and basic metadata associated with a single trait observation. More detailed metadata are available as extensions to this master file, as recommended in the ecological trait-data Standard (Schneider et al. 2019).

***FAIRTraits_Taxon.csv*:** 241 rows (excluding the header); 17 columns; 47 Ko.

***FAIRTraits_EnvironmentalPlots.csv*:** 84 rows (excluding the header); 27 columns; 24 Ko

***FAIRTraits_MeteoSitesMonth.csv*:** 84 rows (excluding the header); 18 columns; 10 Ko

***FAIRTraits_MeteoSitesYear.csv*:** 7 rows (excluding the header); 17 columns; 2 Ko

#### 3. Format and storage mode

All files are provided in tab-separated-value format (csv), encoded with UTF_8.

#### 4. Header information

See variable information (Table 4 in section Class IV.B. below).

#### 5. Special characters/fields

In all files, “NA” denotes missing data, while “none” means that there is no relevant value for the attribute (for example, a simple leaf will have a “none” value for the trait “number of leaflets”).

Three identifiers were designed for each record: the first one is the unique identifier of the record (verbatimOccurrenceID; required for Darwin Core archives: cf. Table 4 in section Class IV.B.1.), while the other two have been designed to enable users to associate data from different traits at the replicate (verbatimOccurrenceID_sample) and population levels (verbatimOccurrenceID_population).

### B. Variable information

#### 1. Taxa

Field identification of taxa was based on local (mostly de Bolòs et al. 1993, Bernard 2008) or national (Coste 1937) floras. Species names (“acceptedNameUsage” in Darwin Core: Table 6) were homogenized using the reference flora for the French Mediterranean region (Tison et al. 2014), while the scientific name (“scientificName” in Darwin Core: Table 6) was taken from the TaxRef (version 16) referential (https://taxref.mnhn.fr/taxref-web/accueil), the most comprehensive taxonomic resource for France (which also covered the 45 species studied in Catalonia). 224 taxa were identified at the species level, 11 at the subspecies level, and 5 at the genus level. The taxon table (described in Table 4) also gives information on life cycle, life form *sensu* Raunkiær (1934), reproductive phenology and geographic distribution of the taxa (all taken from Tison et al. 2014), and on their photosynthetic pathway (mostly derived from the original ^13^C/^12^C isotopic fraction values available in the database, with additional information given by Colin Osborne). We also provide an original successional stage indicator value for 66 of the species, based on Braun-Blanquet et al. (1952), Escarré et al. (1983) and M. Debussche (pers. com.). Darwin Core terms and classes resulting from the mapping of the variables included in the FAIRTraits_Taxon.csv file are also given in Table 4, as this file is a Darwin Core “Checklist” archive on the GBIF infrastructure.

#### 2. The TraitValues core file

Table 5 describes columns of the core file (FAIRTraits_TraitValues.csv), which contains trait data and contextual information about the collection of these data (e.g. location identification and coordinates, treatments, date, methods, projects, etc.), trait semantics (entity, quality, terminological resources, etc.), and the three identifiers described above. Semantics mostly follow Garnier et al. (2017) for above-ground traits, Freschet et al. (2021) for belowground traits and Ely et al. (2021) for leaf gas exchange. Original definitions were coined when necessary. The two last columns of Table 4 show the Darwin Core terms and classes resulting from the mapping with the original names of variables in the FAIRTraits files, a process required to build the Darwin Core “Occurrence” archive on the GBIF infrastructure.

**Table 5.** Structure of the core file FAIRTraits_TraitValues.csv. The Darwin Core terms and classes are given for each variable (the envPlot variable could not be mapped to the Darwin Core standard).

| Name of variable in raw data file | Variable definition | Darwin Core term | Darwin Core class |
| --- | --- | --- | --- |
| SpeciesName | Name of species taken from a local flora (latin binomial without authority) | acceptedNameUsage | Taxon |
| CodeSp | Identification code of the species (4 first letters of genus name and 4 first letters of species name) | taxonID | Taxon |
| Family | The botanical family of the species | family | Taxon |
| LifeForm1 | Primary life form of species *sensu* Raunkiaer (1934) | taxonRemarks | Taxon |
| Site | Name of site | locality | Location |
| Treatment | Experimental treatment applied in the plot, if any | occurrenceRemarks | Occurrence |
| Year | Year of data collection | year | Event |
| Month | Month of data collection | month | Event |
| Day | Day of data collection | day | Event |
| traitName | Complete name of the trait | measurementTraitName | Trait measurement score (v20140515) |
| verbatimTraitName | Abbreviation used for the trait | measurementTraitID | Trait measurement score (v20140515) |
| replicateID | Identification of the replicate measurement | materialSampleID | MaterialSample |
| verbatimTraitValue | Value taken by the trait for the specified replicate measurement | measurementValue | Trait measurement score (v20140515) |
| verbatimTraitUnit | Unit in which the trait value is expressed | measurementUnit | Trait measurement score (v20140515) |
| traitEntityQuality | Formal combination of trait entity and quality (entity_quality) | measurementTraitRemarks | Trait measurement score (v20140515) |
| traitCategory | The category to which the trait is assigned (building on McCormack et al. 2017) | measurementTraitCategory | Trait descriptor (v20140515) |
| variableType | The type of variable: a trait, a condition or a statistic | measurementType | Trait descriptor (v20140515) |
| termSource | Terminological resource in which trait formalism and definition were taken, when available | measurementTraitSource | Trait descriptor (v201405015) |
| basisOfRecord | The specific nature of the data record | basisOfRecord | Record-level |
| traitID | URL or URI of the web resource used for trait formalism and definition, or definition when not available | measurementTraitIdentifier | Trait measurement score (v20140515) |
| samplingProtocol | Short description of number of, and conditions in which, samples were taken for a specific measurement | samplingProtocol | Event |
| measurementMethod | Short description of, or reference to, the method used to determine the measurement or characteristics | measurementMethod | Trait measurement score (v20140515) |
| experimentalBlock | A set of experimental units grouped on the basis of their similarity for one or more variables | locationRemarks | Location |
| traitPlot | Label of the plot where trait(s) were determined | locationID | Location |
| envPlot | Plot associated with the traitPlot, in which environmental data were assessed | not mapped | - |
| countryCode | Standard 2-letter code of the country in which the sites and plots are located (ISO 3166-1-alpha-2) | countryCode | Location |
| plotLatitude | Latitude of the plot (decimal degree) | decimalLatitude | Location |
| plotLongitude | Longitude of the plot (decimal degree) | decimalLongitude | Location |
| plotAltitude | Altitude of the plot (meters above sea level) | verbatimElevation | Location |
| nameOfProject | Name of project during which data were collected (cf. Table 2) | eventRemarks | Event |
| measurementDeterminedBy | Name of the contact person responsible for the acquisition of the data | recordedBy | Occurrence |
| verbatimOccurrenceID | Unique identifier of the data record | occurrenceID | Occurrence |
| verbatimOccurrenceID_sample | Identifier of the replicate on which a specific data record was taken | organismID | Organism |
| verbatimOccurrenceID_population | Identifier of the population (species x site x plot x treatment x date) in which the measured individual was sampled | eventID | Event |

#### 3. Traits

The list of traits grouped by categories, together with the entities on which these were measured is given in Table 6, and a trait map (cf. McCormarck et al. 2017) organized by trait categories and entities is shown on Figure 3.

**Table 6.** List of traits with their abbreviations, and the entities on which these have been measured (entity abbreviation in brackets), and the category to which each trait has been assigned. Traits are sorted by category. Methods, references and units for measurements are given in the csv files. Trait categories build on McCormack et al. (2017).

| Trait category | Trait name | Trait abbreviation | Entity (abbreviation) |
| --- | --- | --- | --- |
| Allocation ratio | Length fraction of roots with diameter ]0.2-1mm] | RLF]0.2-1mm] | absorptive (abr) or whole root system (wrs) |
| Allocation ratio | Length fraction of roots with diameter ]0-0.2mm] | RLF]0-0.2mm] | absorptive (abr) or whole root system (wrs) |
| Allocation ratio | Length fraction of roots with diameter ]1-2mm] | RLF]1-2mm] | absorptive roots (abr) |
| Allocation ratio | Leaf mass fraction | LMF | whole plant (wpl) |
| Allocation ratio | Absorptive root mass fraction | RabrMF | whole plant (wpl) |
| Allocation ratio | Reproductive dry mass fraction | RepMF | whole plant (wpl) |
| Allocation ratio | Transport root mass fraction | RtrrMF | whole plant (wpl) |
| Allocation ratio | Whole root system mass fraction | RwrsMF | whole plant (wpl) |
| Allocation ratio | Stem mass fraction | StMF | whole plant (wpl) |
| Allocation ratio | Length fraction of roots with diameter higher than 1 mm | RLF>1mm | whole root system (wrs) |
| Architecture | Altitude of the whole root system | RAlt | whole root system (wrs) |
| Architecture | Magnitude of the whole root system | RMag | whole root system (wrs) |
| Architecture | Root topological index | RTopo | whole root system (wrs) |
| Architecture | Number of root tips per unit of root length of the whole root system | TipNbRL | whole root system (wrs) |
| Chemistry | Root nitrogen content per root dry mass | RNC | absorptive (abr) or whole root system (wrs) |
| Chemistry | Root carbon content per root dry mass | RCC | absorptive (abr), transport (trr) or whole root system (wrs) |
| Chemistry | Root cellulose content per root dry mass | RCellulose | absorptive roots (abr) |
| Chemistry | Root carbon isotopic fraction | Rdelta13C | absorptive roots (abr) |
| Chemistry | Root nitrogen isotopic fraction | Rdelta15N | absorptive roots (abr) |
| Chemistry | Root hemicellulose content per root dry mass | RHemicell | absorptive roots (abr) |
| Chemistry | Root lignin content per root dry mass | RLignin | absorptive roots (abr) |
| Chemistry | Root water-soluble compounds content per root dry mass | RSoluble | absorptive roots (abr) |
| Chemistry | Leaf aluminum content per leaf dry mass | LAl | mature leaf (mlf) |
| Chemistry | Leaf boron content per leaf dry mass | LB | mature leaf (mlf) |
| Chemistry | Leaf calcium content per leaf dry mass | LCa | mature leaf (mlf) |
| Chemistry | Leaf chromium content per leaf dry mass | LCr | mature leaf (mlf) |
| Chemistry | Leaf copper content per leaf dry mass | LCu | mature leaf (mlf) |
| Chemistry | Leaf nitrogen isotopic fraction | Ldelta15N | mature leaf (mlf) |
| Chemistry | Leaf oxygen isotopic fraction | Ldelta18O | mature leaf (mlf) |
| Chemistry | Leaf iron content per leaf dry mass | LFe | mature leaf (mlf) |
| Chemistry | Leaf potassium content per leaf dry mass | LK | mature leaf (mlf) |
| Chemistry | Leaf magnesium content per leaf dry mass | LMg | mature leaf (mlf) |
| Chemistry | Leaf manganese content per leaf dry mass | LMn | mature leaf (mlf) |
| Chemistry | Leaf sodium content per leaf dry mass | LNa | mature leaf (mlf) |
| Chemistry | Leaf nickel content per leaf dry mass | LNi | mature leaf (mlf) |
| Chemistry | Leaf lead content per leaf dry mass | LPb | mature leaf (mlf) |
| Chemistry | Leaf sulfur content per leaf dry mass | LS | mature leaf (mlf) |
| Chemistry | Leaf strontium content per leaf dry mass | LSr | mature leaf (mlf) |
| Chemistry | Leaf titanium content per leaf dry mass | LTi | mature leaf (mlf) |
| Chemistry | Leaf zinc content per leaf dry mass | LZn | mature leaf (mlf) |
| Chemistry | Leaf carbon content per leaf dry mass | LCC | mature leaf (mlf), rep. (rhs), veg. (vsh) or whole shoot (wsh) |
| Chemistry | Leaf carbon isotopic fraction | Ldelta13C | mature leaf (mlf), rep. (rhs), veg. (vsh) or whole shoot (wsh) |
| Chemistry | Leaf nitrogen content per leaf dry mass | LNC | mature leaf (mlf), rep. (rhs), veg. (vsh) or whole shoot (wsh) |
| Chemistry | Leaf phosphorus content per leaf dry mass | LPC | mature leaf (mlf), rep. (rhs), veg. (vsh) or whole shoot (wsh) |
| Chemistry | Reproductive organ carbon content per reproductive organ dry mass | RepCC | reproductive system (rep) |
| Chemistry | Reproductive organ nitrogen content per reproductive organ dry mass | RepNC | reproductive system (rep) |
| Chemistry | Stem carbon content per leaf dry mass for graminoid species | StCC | sheath (she) |
| Chemistry | Stem nitrogen content per leaf dry mass for graminoid species | StNC | sheath (she) |
| Chemistry | Stem carbon content per leaf dry mass for non-graminoid species | StCC | stem (ste) |
| Chemistry | Stem nitrogen content per leaf dry mass for non-graminoid species | StNC | stem (ste) |
| Chemistry | Dead shoot carbon content per dead shoot dry mass | ShDeadCC | vegetative (vsh) or reproductive (rsh) shoot |
| Chemistry | Dead shoot nitrogen content per dead shoot dry mass | ShDeadNC | vegetative (vsh) or reproductive (rsh) shoot |
| Chemistry | Stem (or sheath) carbon content per stem (or sheath) dry mass | StCC | vegetative (vsh), reproductive (rsh) or whole shoot (wsh) |
| Chemistry | Stem (or sheath) nitrogen content per stem (or sheath) dry mass | StNC | vegetative (vsh), reproductive (rsh) or whole shoot (wsh) |
| Chemistry | Stem (or sheath) phosphorus content per stem (or sheath) dry mass | StPC | vegetative (vsh), reproductive (rsh) or whole shoot (wsh) |
| Chemistry | Shoot carbon content per shoot dry mass | ShCC | vegetative shoot (vsh) |
| Chemistry | Shoot nitrogen content per shoot dry mass | ShNC | vegetative shoot (vsh) |
| Chemistry | Root phosphorus content per root dry mass | RPC | whole root system (wrs) |
| Condition | CO<sub>2</sub> concentration in wet air inside chamber | CO<sub>2</sub>s | chamber condition (chc) |
| Condition | Atmospheric pressure | P<sub>atm</sub> | chamber condition (chc) |
| Condition | In-chamber photosynthetic flux density (PPFD) incident on the leaf, quanta per area | Q<sub>in</sub> | chamber condition (chc) |
| Condition | Relative humidity of air inside the chamber | RHs | chamber condition (chc) |
| Condition | Time of day for gas exchange measurement | timeOfDay | chamber condition (chc) |
| Dynamics | Root productivity | RProd | absorptive roots (abr) |
| Dynamics | Root remaining mass | RRM84 | absorptive roots (abr) |
| Dynamics | Leaf litter remaining mass | LLitRM56 | leaf litter (llt) |
| Dynamics | Leaf litter remaining mass calculated | LLitRM56c | leaf litter (llt) |
| Dynamics | Stem litter remaining mass | StLitRM56 | stem litter (stl) |
| Mechanics | Leaf tensile strength | LTS | mature leaf (mlf) |
| Mechanics | Leaf work to shear | LWS | mature leaf (mlf) |
| Microbial associations | Root mycorrhizal colonization frequency | FqMyc | absorptive roots (abr) |
| Microbial associations | Root mycorrhizal colonization intensity | InMyc | absorptive roots (abr) |
| Morphology | Root tissue density | RTD | absorptive (abr) or whole root system (wrs) |
| Morphology | Average root diameter | RD | absorptive (abr), transport (trr) or whole root system (wrs) |
| Morphology | Specific root area | SRA | absorptive (abr), transport (trr) or whole root system (wrs) |
| Morphology | Specific root length | SRL | absorptive (abr), transport (trr) or whole root system (wrs) |
| Morphology | Root dry matter content | RDMC | absorptive roots (abr) |
| Morphology | Root dry matter content of deep absorptive roots | RDMC | deep absorptive roots (dar) |
| Morphology | Root tissue density of deep absorptive roots | RTD | deep absorptive roots (dar) |
| Morphology | Specific root length of deep absorptive roots | SRL | deep absorptive roots (dar) |
| Morphology | Fruit dry mass | FtDM | fruit (frt) |
| Morphology | Area of a leaf | LArea | mature leaf (mlf) |
| Morphology | Dry mass of a leaf | LDM | mature leaf (mlf) |
| Morphology | Leaflet width | LfletWidth | mature leaf (mlf) |
| Morphology | Fresh mass of a leaf | LFM | mature leaf (mlf) |
| Morphology | Leaf length | LLength | mature leaf (mlf) |
| Morphology | Leaf lamina thickness excluding midvein | LT | mature leaf (mlf) |
| Morphology | Leaf width | LWidth | mature leaf (mlf) |
| Morphology | Specific leaf area | SLA | mature leaf (mlf) |
| Morphology | Leaf dry matter content | LDMC | mature leaf (mlf) or whole shoot (wsh) |
| Morphology | Seed depth | SdDepth | mature seed (msd) |
| Morphology | Seed dry mass | SdDM | mature seed (msd) |
| Morphology | Seed length | SdLength | mature seed (msd) |
| Morphology | Seed width | SdWidth | mature seed (msd) |
| Morphology | Stem dry matter content for graminoid species | StDMC | sheath (she) |
| Morphology | Stem dry matter content for non-graminoid species | StDMC | stem (ste) |
| Morphology | Root dry matter content | RDMC | whole root system (wrs) |
| Morphology | Stem (or sheath) dry matter content | StDMC | whole shoot (wsh) |
| Phenology | Leaf life span | LLS | leaf cohort (clf) |
| Phenology | Time of seed dispersal | Disp | reproductive system (rep) |
| Phenology | Onset of flowering | Flo | reproductive system (rep) |
| Phenology | Duration of seed maturation period | MatPer | reproductive system (rep) |
| Physiology | Root specific respiration | RSR | absorptive roots (abr) |
| Physiology | Net CO<sub>2</sub> exchange per leaf area | A<sub>area</sub> | mature leaf (mlf) |
| Physiology | Net CO<sub>2</sub> exchange per leaf dry mass | A<sub>mass</sub> | mature leaf (mlf) |
| Physiology | Intercellular CO<sub>2</sub> concentration in air | Ci | mature leaf (mlf) |
| Physiology | Transpiration rate of H<sub>2</sub>O per leaf area | E<sub>area</sub> | mature leaf (mlf) |
| Physiology | Stomatal conductance to water vapor per leaf area | g<sub>sw</sub> | mature leaf (mlf) |
| Physiology | Leaf surface temperature | Tleaf | mature leaf (mlf) |
| Plant size | Root mass density | RMD | absorptive (abr) or whole root system (wrs) |
| Plant size | Root length density | RLD | absorptive (abr), transport (trr) or whole root system (wrs) |
| Plant size | Total leaf dry mass of the plant | LDM | mature leaf (mlf), rep. (rsh), veg. (vsh) or whole shoot (wsh) |
| Plant size | Maximum horizontal plant length when plant in reproductive stage | Dmax | reproductive shoot (rsh) |
| Plant size | Maximum horizontal plant width when plant in reproductive stage | Dmin | reproductive shoot (rsh) |
| Plant size | Reproductive plant height | Hrep | reproductive shoot (rsh) |
| Plant size | Total number of seeds produced by the plant | SdNb | reproductive shoot (rsh) |
| Plant size | Total dry mass of reproductive organs | RepDM | reproductive system (rep) |
| Plant size | Total dry mass of reproductive shoots | RepDM | reproductive system (rep) |
| Plant size | Total fresh mass of reproductive organs | RepFM | reproductive system (rep) |
| Plant size | Bulb dry mass | BDM | stem (ste) |
| Plant size | Bulb fresh mass | BFM | stem (ste) |
| Plant size | Dead shoot dry mass | ShDeadDM | vegetative (vsh), reproductive (rsh) or whole shoot (wsh) |
| Plant size | Shoot dry mass | ShDM | vegetative (vsh), reproductive (rsh) or whole shoot (wsh) |
| Plant size | Total stem (or sheath) dry mass of the plant | StDM | vegetative (vsh), reproductive (rsh) or whole shoot (wsh) |
| Plant size | Maximum horizontal plant length when plant in vegetative stage | Dmax | vegetative shoot (vsh) |
| Plant size | Maximum horizontal plant width when plant in vegetative stage | Dmin | vegetative shoot (vsh) |
| Plant size | Vegetative plant height | Hveg | vegetative shoot (vsh) |
| Plant size | Total dry mass of the whole plant | WPDM | whole plant (wpl) |
| Plant size | Rooting depth | RDepth | whole root system (wrs) |
| Plant size | Rooting depth 95% | RDepth95 | whole root system (wrs) |
| Plant size | Root dry mass | RDM | whole root system (wrs) |
| Plant size | Root fresh mass | RFM | whole root system (wrs) |
| Plant size | Maximum length of the stretched root system | RLStretch | whole root system (wrs) |
| Plant size | Total dead leaf dry mass of the plant | DLDM | whole shoot (wsh) |
| Plant size | Total dead leaf fresh mass of the plant | DLFM | whole shoot (wsh) |
| Plant size | Total leaf fresh mass of the plant | LFM | whole shoot (wsh) |
| Plant size | Total stem (or sheath) fresh mass of the plant | StFM | whole shoot (wsh) |
| Statistics | Higher confidence limit of calculated leaf litter remaining mass | HiLLitRM56c | leaf litter (llt) |
| Statistics | Lower confidence limit of calculated leaf litter remaining mass | LoLLitRM56c | leaf litter (llt) |

**Figure 3.**
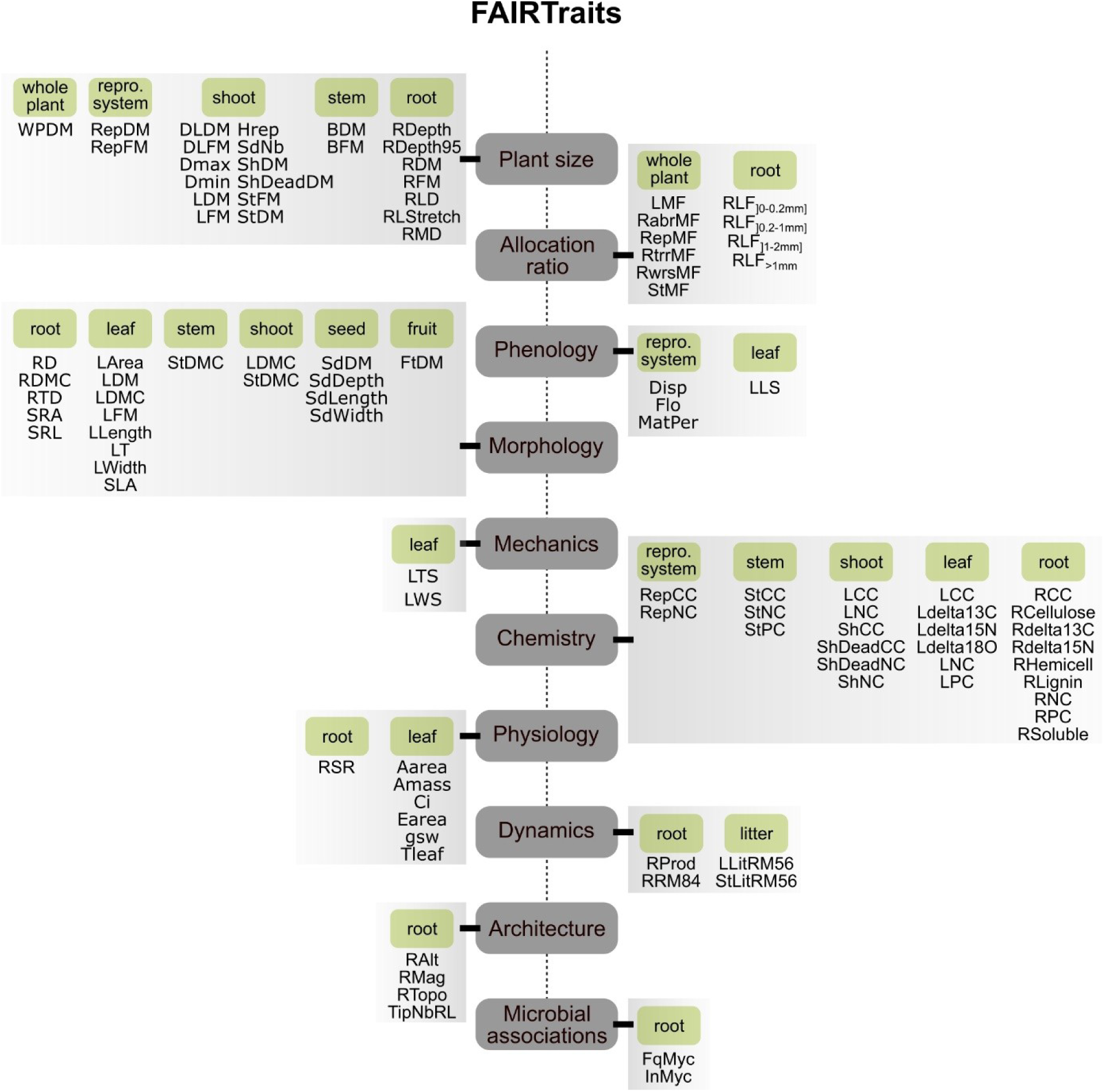
Map of the documented traits documented in the FAIRTraits database. The traits are grouped into 10 trait categories (allocation ratio, architecture, chemistry, dynamics, mechanics, microbial associations, morphology, phenology, physiology, plant size) and 9 primary entities (whole plant, reproductive system, fruit, seed, shoot, stem, leaf, litter, root). For the sake of readibility, the most significant traits to each category and entity are shown on the map. See Table 6 for trait abbreviations, details on secondary entities, and for the complete list of traits.

Data for a total number of 183 traits are available in the database (Table 6). Each trait corresponds to a unique quality x entity combination, which is the formalism followed to describe traits (Mungall et al. 2010, Garnier et al. 2017). Overall, qualities were measured on 17 entities, detailed in Table 6. For example, root nitrogen content per root dry mass (RNC; a quality) has been measured on three entities: absorptive roots (abr), transport roots (trr) and the whole root system (wrs). There are thus three quality x entity combinations for this specific quality, i.e. three traits, which are distinguished in the database: RNC_abr, RNC_trr and RNC_wrs, respectively. Size-related traits (plant height, horizontal dimensions) measured on shoots are the most represented traits in the database, followed by traits pertaining to leaf morpho-anatomy and chemical composition, including leaf carbon isotopic ratio (Figure 4d). Root and seed traits constitute a substantial part of the records as well (Figure 4e). We also provide data on several physiological traits: leaf gas exchange (net photosynthesis, transpiration, intrinsic water-use efficiency) for 60 species (70 populations), root respiration for 16 species, leaf litter and root decomposition for 42 species and 33 species, respectively. Flowering onset and time of seed dispersal (reproductive phenology) are available for 160 and 164 species, respectively.

**Figure 4.**
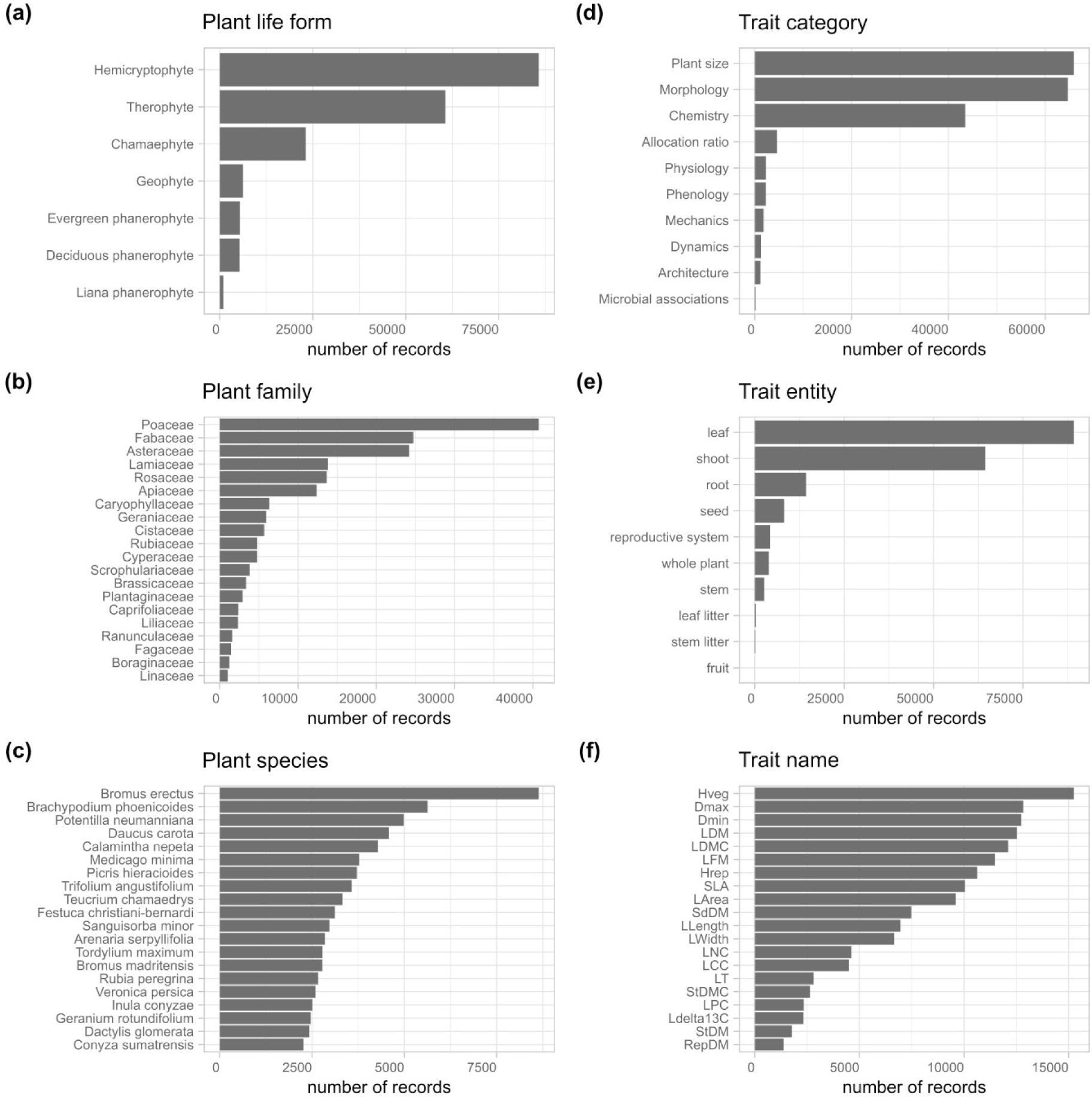
Number of records in the FAIRTraits database organized by (a) plant life form *sensu* Raunkiær (1934), (b) botanical family for the 20 most represented families, (c) species for the 20 most represented species, (d) trait category, (e) trait entity and (f) trait name for the 20 most documented traits. See Table 5 for trait abbreviations.

The number of records, minimum, maximum and distribution of trait values together with their unit are given in Figure 5.

**Figure 5.**
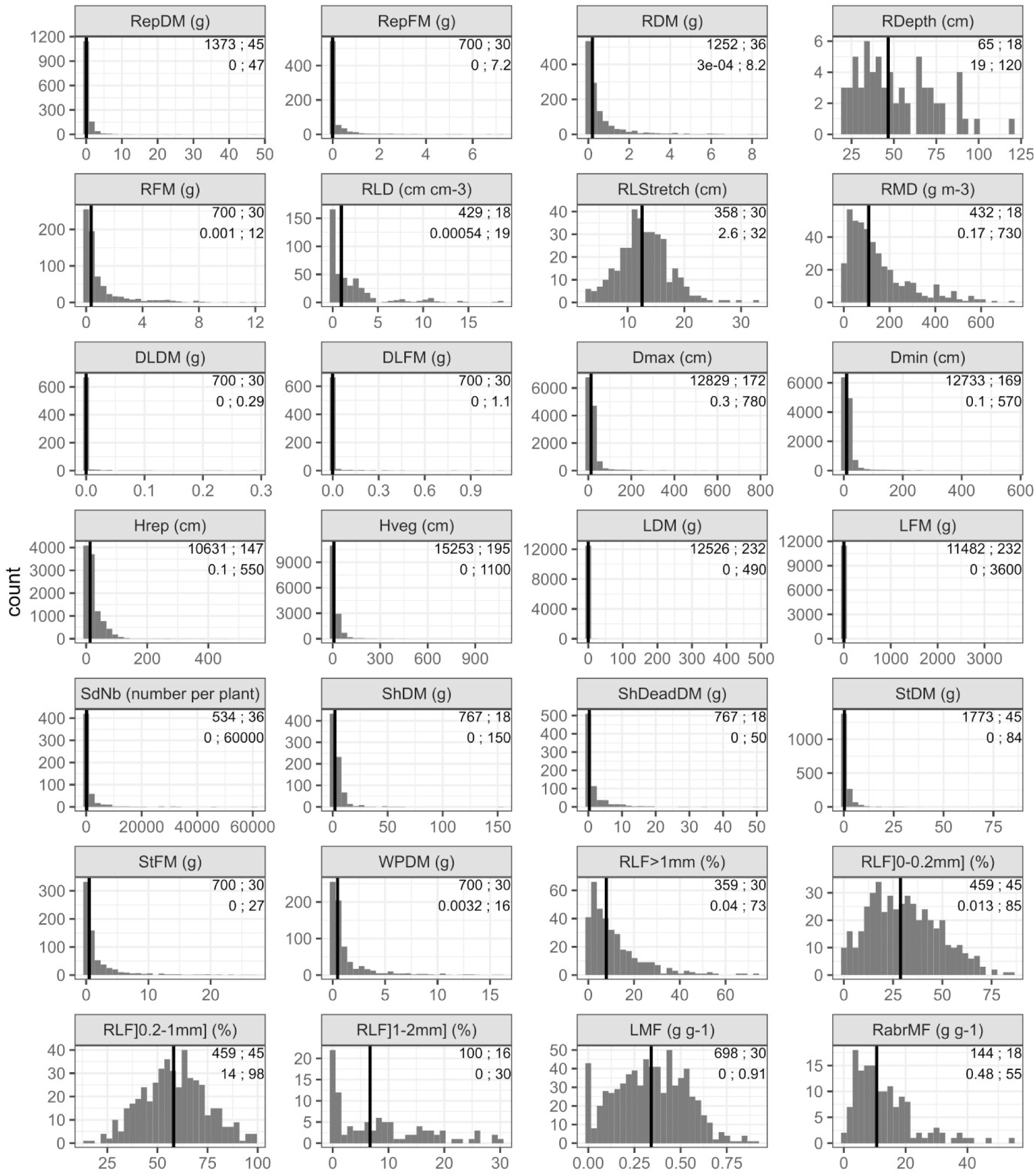
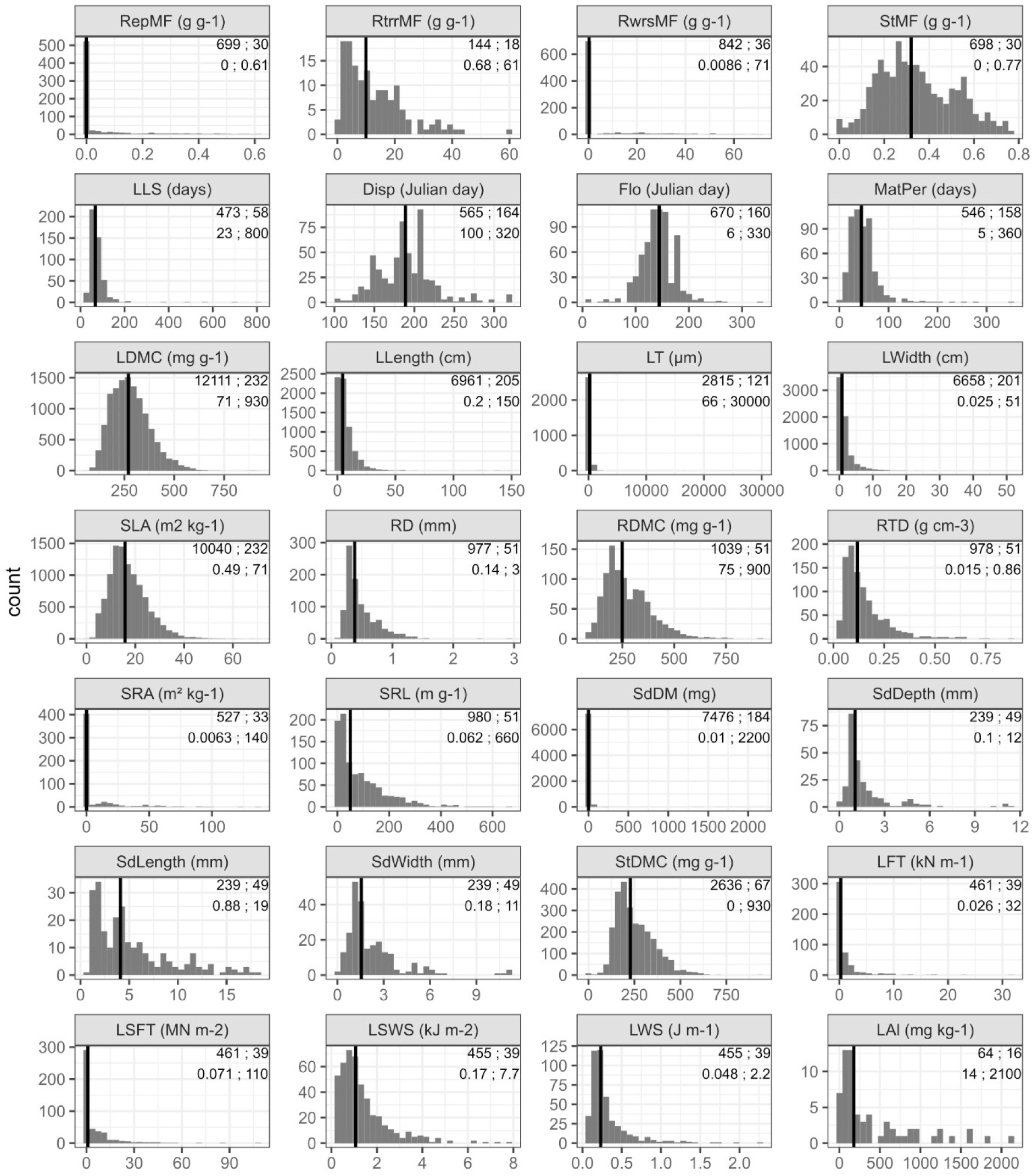
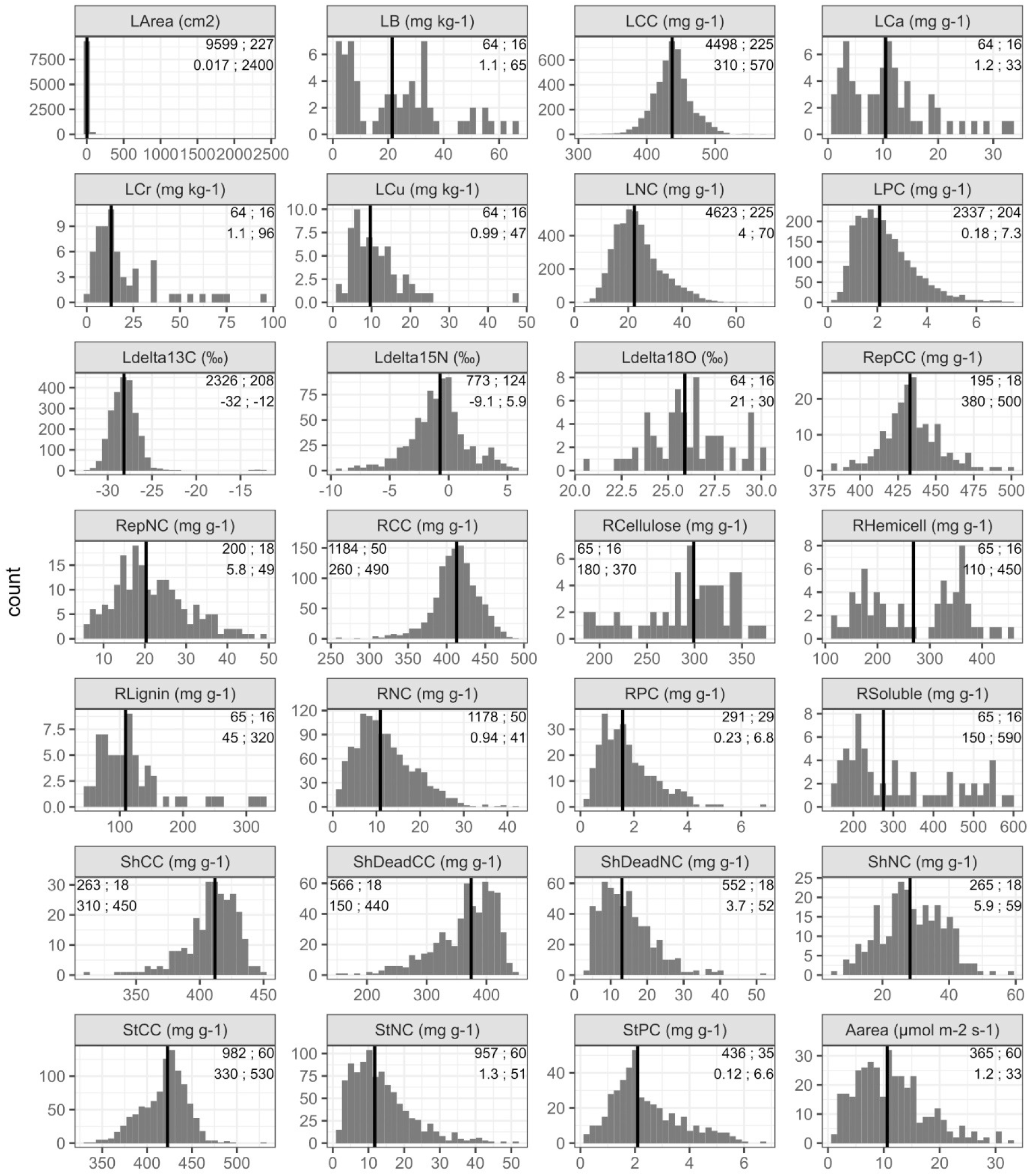
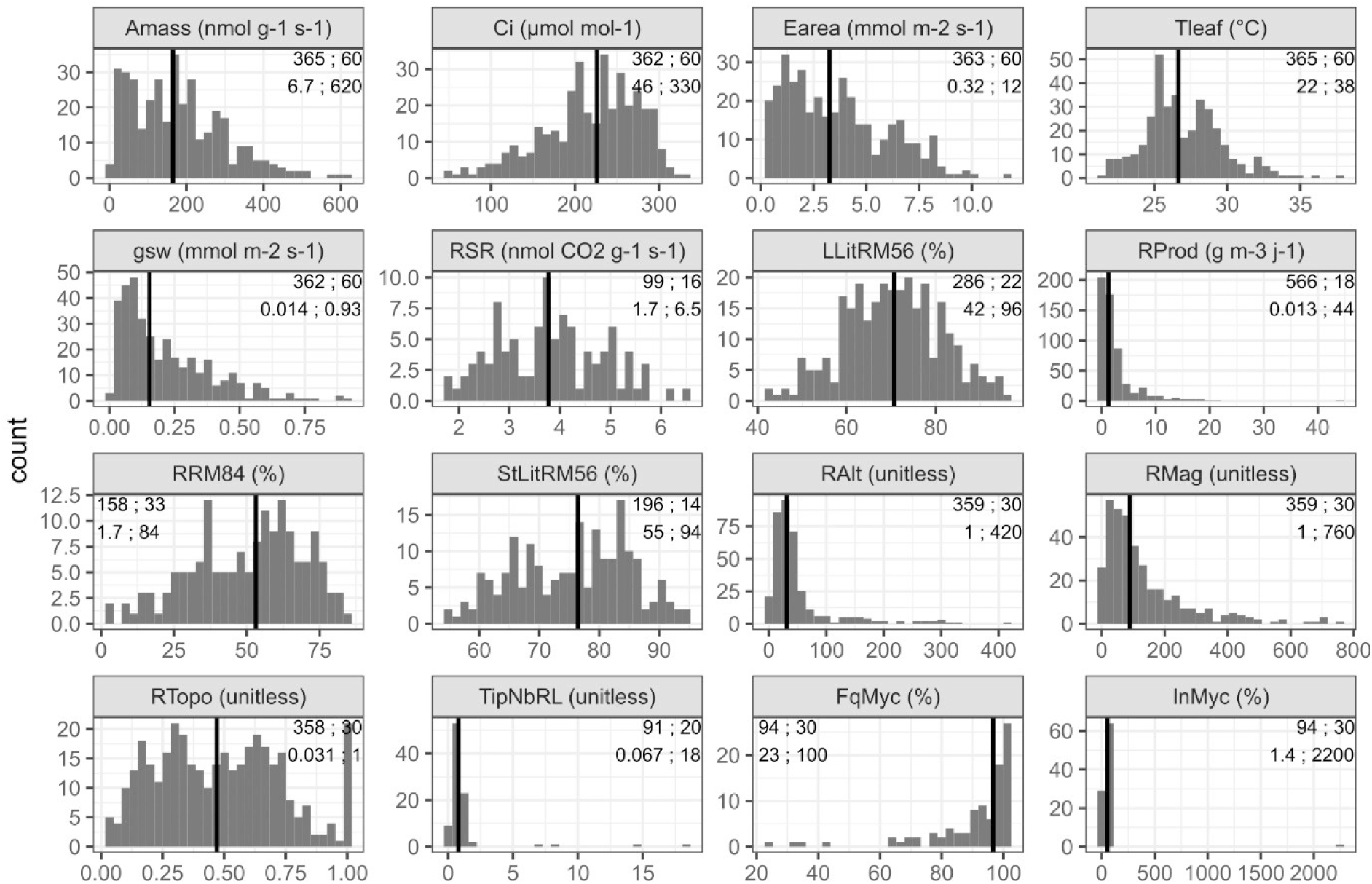
Frequency distributions of trait values. Units are shown next to the trait abbreviations (Table 5 for full names). Figures inside each panel show the number of records and number of species (upper line), and minimum and maximum values (lower line). On each panel, trait values for all entities corresponding to a given quality are shown.

#### 4. Environmental data

Trait values not only depend on the taxa on which they are determined, but also on the environmental conditions in which the individuals representing these taxa are sampled (Violle et al. 2007, Garnier et al. 2016, Shipley et al. 2016, Anderegg 2023 for references and discussions). To describe the environmental context in which the FAIRTraits data were collected, we provide information on climatic variables at the site level taken from the meteorological station closest to each site (cf. Table 3 and Figure 1 for climatic diagrams, in section Class II.B.1.), and on treatments, disturbance (type, intensity and return interval) and soil characteristics at the plot level (see section Class III.B.2.a. above for details on the design). The list of environmental variables available in the database is given in Table 7. Since most of these environmental variables are not defined in the Darwin Core standard, these data are not available as a Darwin Core archive. They are archived as csv files in the InDoRES repository.

**Table 7.** List of variables in the three files describing environmental data: FAIRTraits_EnvironmentalPlots.csv (envPlot), FAIRTraits_MeteoSitesYear.csv (annMet) and FAIRTraits_MeteoSitesMonth.csv files (monMet) files. Methods, references and units are given in the csv files.

| Name of variable | File | Name of variable |
| --- | --- | --- |
| siteName | monMet, annMet & envPlot | Name of site |
| siteLatitude | monMet & annMet | Latitude of site |
| siteLongitude | monMet & annMet | Longitude of site |
| siteMinAltitude | monMet & annMet | Minimum altitude of site |
| siteMaxAltitude | monMet & annMet | Maximum altitude of site |
| siteMeteoStationName | monMet & annMet | Name of meteorological station for site |
| siteMeteoStationID | monMet & | Identifier of meteorological station for site |
|  | annMet |  |
| siteDistMeteoStation | monMet & annMet | Distance from site to meteorological station |
| siteMeteoStationLatitude | monMet & annMet | Latitude of meteorological station for site |
| siteMeteoStationLongitude | monMet & annMet | Longitude of meteorological station for site |
| siteMeteoStationAltitude | monMet & annMet | Altitude of meteorological station for site |
| siteMeteoStationCalculationPeriod | monMet & annMet | Calculation period for meteorological data |
| siteMeteoStationMAT | annMet | Mean annual temperature from meteorological station |
| siteMeteoStationMColdM | annMet | Mean temperature of the coldest month from meteorological station |
| siteMeteoStationMWarM | annMet | Mean temperature of the warmest month from meteorological station |
| siteMeteoStationMAP | annMet | Mean annual precipitation from meteorological station |
| siteMeteoStationETP | annMet | Mean annual potential evaporation from meteorological station |
| siteMeteoStationMonth | monMet | Month of data record from the meteorological station |
| siteMeteoStationMMT | monMet | Mean monthly temperature from meteorological station |
| siteMeteoStationMMMAXT | monMet | Mean monthly maximal temperature from meteorological station |
| siteMeteoStationMMMINT | monMet | Mean monthly minimal temperature from meteorological station |
| siteMeteoStationMMP | monMet | Mean monthly precipitation from meteorological station |
| siteMeteoStationMMETP | monMet | Mean monthly potential evaporation from meteorological station |
| envPlotID | envPlot | Plot identification code |
| envPlotLatitude | envPlot | Latitude of environmental plot |
| envPlotLongitude | envPlot | Longitude of environmental plot |
| envPlotAltitude | envPlot | Altitude of environmental plot |
| treatment | envPlot | Identification of treatment |
| treatmentDescription | envPlot | Short description of treatment |
| refTreatmentDescription | envPlot | Literature reference for detailed description of treatment |
| envPlotDisturbanceType | envPlot | Type of disturbance in plot (grazing, cultivation, etc.) |
| envPlotDisturbanceIntensity | envPlot | Disturbance intensity in plot |
| envPlotDisturbanceReturnInterval | envPlot | Return time interval of disturbance in plot |
| envPlotFertilization | envPlot | Level of fertilization in plot |
| envPlotSoilType | envPlot | Type of soil in plot |
| envPlotPeriodSoilAnalyses | envPlot | Time when soil samples were collected for analyses |
| envPlotSoilLayerAnalyzed | envPlot | Minimum and maximum depths of soil sampled for analyses |
| envPlotHumidity | envPlot | Humidity of fresh soil |
| envPlotClay | envPlot | Clay fraction of soil (particle size < 2 µm) |
| envPlotSilt | envPlot | Silt fraction of soil (particle size between 2 and 50 µm) |
| envPlotSand | envPlot | Sand fraction of soil (particle size > 50 µm) |
| envPlotCaCO3 | envPlot | Total lime content of soil per mass |
| envPlotCorg | envPlot | Organic carbon content of soil per mass |
| envPlotNTotal | envPlot | Total (organic + mineral) nitrogen content of soil per mass |
| envPlotCNRatio | envPlot | Ratio of organic content to total nitrogen content of soil |
| envPlotOrgMat | envPlot | Organic matter content of soil per unit mass |
| envPlotpH | envPlot | pH in water |
| envPlotPOlsen | envPlot | Extractable phosphorus content of soil per mass |
| envPlotCEC | envPlot | Cation-exchange capacity of soil per mass |
| aggregationMethod | envPlot | Method used to infer soil values for plots with unavailable data |

## Class V. Supplemental descriptors

### A. Data acquisition

#### 1. Location of completed data forms

Paper data sheets still available from the original field data collections are stored by the ECOPAR research group in the premises of the Centre d’Écologie Fonctionnelle et Évolutive (CNRS Campus: 1919 route de Mende, 34293 Montpellier Cedex, France).

#### 2. Data entry verification procedures

Field and laboratory data were first digitized by the person(s) in charge of the observations and measurements in spreadsheet files of different formats. Subsets of data were checked and cleaned prior to analyses conducted in the context of the different research projects. Data on reproductive phenology was first consolidated across the whole database by Jules Segrestin (see Segrestin et al. 2018). The current version of FAIRTraits was compiled and reviewed by Eric Garnier, Léo Delalandre, Karim Barkaoui and Jules Segrestin.

### B. Quality assurance/quality control procedures

The database underwent extensive quality assurance and control processes before publication. Trait names, taxon names, plot inventories, unexpected missing data, and inconsistencies in methods were corrected during the structuring process, which converted native files into the final database. These steps are described in detail in Table 8 (Section Class V.C.).

**Table 8.**
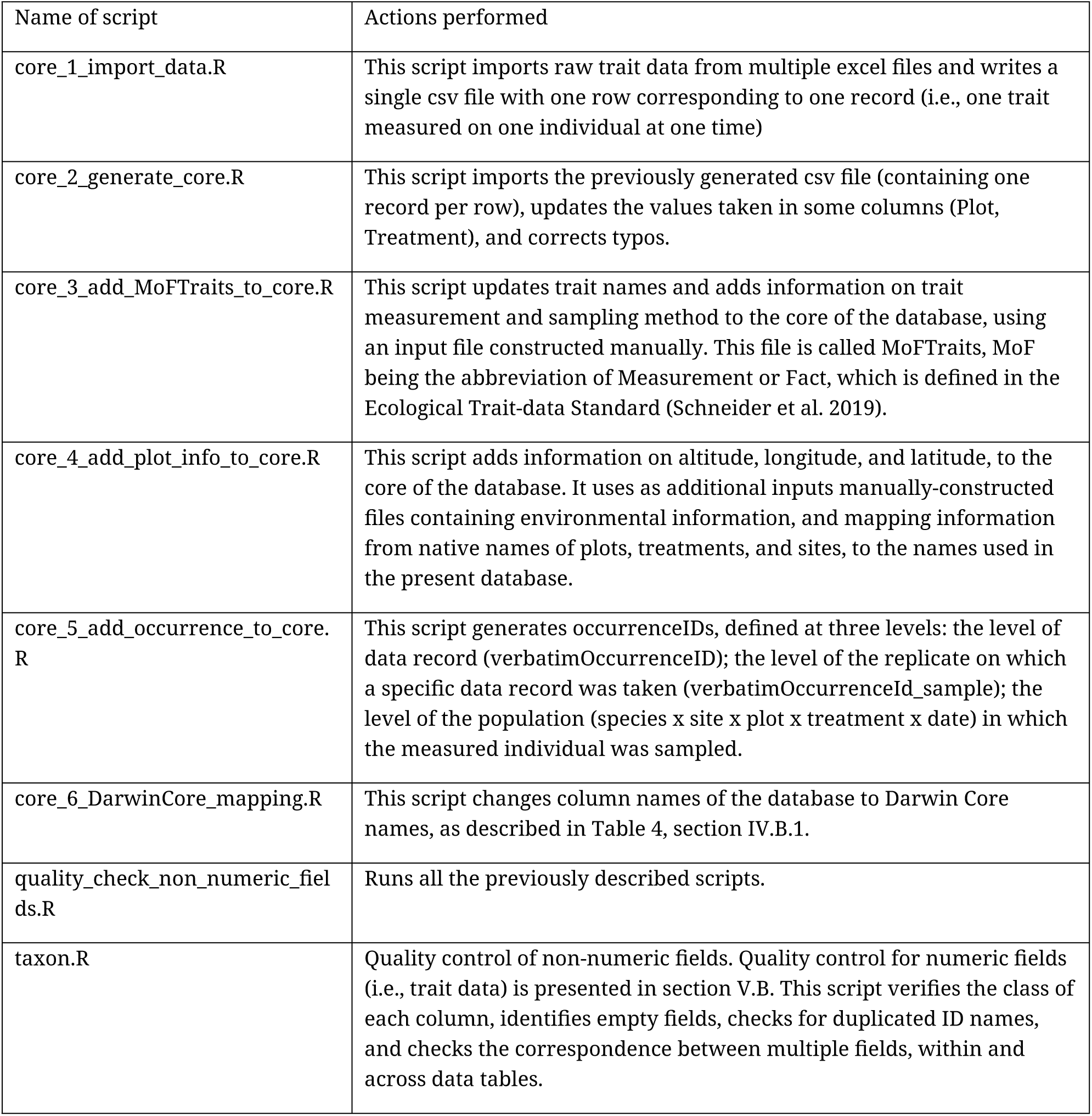
Description of the scripts used to process the data from native format to the present database.

| Name of script | Actions performed |
| --- | --- |
| core_1_import_data.R | This script imports raw trait data from multiple excel files and writes a single csv file with one row corresponding to one record (i.e., one trait measured on one individual at one time) |
| core_2_generate_core.R | This script imports the previously generated csv file (containing one record per row), updates the values taken in some columns (Plot, Treatment), and corrects typos. |
| core_3_add_MoFTraits_to_core.R | This script updates trait names and adds information on trait measurement and sampling method to the core of the database, using an input file constructed manually. This file is called MoFTraits, MoF being the abbreviation of Measurement or Fact, which is defined in the Ecological Trait-data Standard (Schneider et al. 2019). |
| core_4_add_plot_info_to_core.R | This script adds information on altitude, longitude, and latitude, to the core of the database. It uses as additional inputs manually-constructed files containing environmental information, and mapping information from native names of plots, treatments, and sites, to the names used in the present database. |
| core_5_add_occurrence_to_core.R | This script generates occurrenceIDs, defined at three levels: the level of data record (verbatimOccurrenceID); the level of the replicate on which a specific data record was taken (verbatimOccurrenceId_sample); the level of the population (species x site x plot x treatment x date) in which the measured individual was sampled. |
| core_6_DarwinCore_mapping.R | This script changes column names of the database to Darwin Core names, as described in Table 4, section IV.B.1. |
| quality_check_non_numeric_fields.R | Runs all the previously described scripts. |
| taxon.R | Quality control of non-numeric fields. Quality control for numeric fields (i.e., trait data) is presented in section V.B. This script verifies the class of each column, identifies empty fields, checks for duplicated ID names, and checks the correspondence between multiple fields, within and across data tables. |

The final version of FAIRTraits includes values for 181 traits measured across 1 to 232 species. Error detection for each trait and species followed a two-step approach:

First for species with sufficient replicates (n > 4), we tested the consistency of units by assessing whether the distribution of trait values across all replicates followed a unimodal pattern. This was done using the *dip.test* function from the *diptest* R package (Maechler, 2024). For distributions with significant results (P < 0.001), we manually inspected the data and corrected any discrepancies in unit expression. This process identified cases where field campaigns (e.g., a sampling season at one site) systematically recorded a trait using a different unit (e.g., meters instead of centimeters for plant height). In such cases, all replicate trait values for all species recorded during the campaign were corrected accordingly.

Second, errors in individual observations were identified using the Grubbs’ test for outlier detection (*grubbs.test* function in the *outliers* R package; Komsta, 2022). The test was applied separately to each trait and species with at least five replicates. For significant results (P < 0.05), we manually inspected the data. Obvious errors were corrected, while values determined to be true outliers were conservatively removed from the database following recommendations from Keller et al. (2023).

### C. Computer programs and data-processing algorithms

The re-structuring of data from native files to the database presented here was performed using the software R (R Core Team 2020). All the scripts used for processing the data are available on GitHub (https://github.com/LDelalandre/FAIRTraits). They are described in Table 8.

### D. Archiving

#### 1. Archival procedures: (to be finalized)

Metadata are archived in the Cat.InDoRES metadata catalog (https://cat.indores.fr/geonetwork/srv/eng/catalog.search#/home). The part of the database which was possible to format with the Darwin Core standard (see Table 1 in section Class I.B.) is archived in the GBIF infrastructure (https://www.gbif.org/fr/), while the remaining files listed in Table 1 (section Class I.B.) are available in the InDoRES repository (https://www.indores.fr/), The associated R code used to process the files is available on GitHub.

#### 2. Redundant archival sites

The list of doi in different repositories and cross-references among repositories will be described here.

### E. Publications and results

Subsets of the database have been used in studies leading to the following publications (associated research projects are given in the FAIRTraits_PublishedPapers.csv file).

*An “*” in front of the name of the first author indicates that the subset has been combined with data from other origins in broad compilations and/or analyses*.

### F. History of data set usage

1. **Data request history:** see publications with an “*” in front of the name of the first author in section Class V.B. (“Publications and results”) above. For these large compilations and/or analyses, data aggregated at the species x treatment level were provided.
2. **Data set update history:** the version of FAIRTraits as described in this data paper is the first complete version of the database. It includes original data beyond those used in the publications listed in section Class V.B.
3. 

## Acknowledgements

We thank Anabelle Dos Santos, Magdy El-Bana and the numerous students and technicians who contributed to the collection and processing of samples during the successive field campaigns throughout the period covered by the different projects. Many thanks to: James Aronson and Edouard Le Floc’h for creating favourable conditions for research at the Cazarils site; to Joël and Danièle Garnier for the re-birth of Les Agros; to the technical staff of the INRAE experimental farm at La Fage and of the CEFE experimental garden in Montpellier, a technical platform from the CeMEB LabEx (Centre Méditerranéen de l’Environnement et de la Biodiversité; ANR-10-LABX-0004 convention). Max Debussche and Colin Osborne consolidated the assignment of species to successional stages and photosynthetic pathways, respectively; Félix de Tombeur led us through the labyrinth of the WRB soil classification. The research projects which led to the initial collection of data are listed in Table 2 of this metadata document (section Class II.A.6.). This paper has benefited from the most welcome and enthusiastic assistance of Sophie Pamerlon (GBIF France), Yvan Le Bras and Olivier Norvez (both at the Pôle National des Données de la Biodiversité), in the context of the OpenMetaPaper project funded by the French Ministry of Higher Education and Research (FNSO project AAPFNSO2019OpenMetaPaper-14026).

## References

Barkaoui, K., Bernard-Verdier, M., Navas, M.-L. (2013). Questioning the reliability of the point intercept method for assessing community functional structure in low-productive and highly diverse Mediterranean grasslands. Folia Geobotanica, 48 (3): 393–414. 10.1007/s12224-013-9172-2.

Barkaoui, K., Roumet, C., & Volaire, F. (2016). Mean root trait more than root trait diversity determines drought resilience in native and cultivated Mediterranean grass mixtures. Agriculture, Ecosystems & Environment, 231: 122–132. 10.1016/j.agee.2016.06.035.

Bernard-Verdier, M., Flores, O., Navas, M.-L., Garnier E. (2013). Partitioning phylogenetic and functional diversity into alpha and beta components along an environmental gradient in a Mediterranean rangeland. Journal of Vegetation Science, 24: 877–889. 10.1111/jvs.12048.

Bernard-Verdier, M., Navas, M.-L., Vellend, M., Violle, C., Fayolle, A. & Garnier, E. (2012). Community assembly along a soil depth gradient: contrasting patterns of plant trait convergence and divergence in a Mediterranean rangeland. Journal of Ecology, 100: 1422–1433. 10.1111/1365-2745.12003.

Birouste M., Kazakou E. Blanchard A., Roumet C. 2012. Plants traits and decomposition: are the relationships for roots comparable to those for leaves? Annals of Botany, 109: 463–472. 10.1093/aob/mcr297.

Bumb, I., Garnier, E., Bastianelli, D., Richarte, J., Bonnal, L., Kazakou, E. (2016). Influence of management regime and harvest date on the forage quality of rangelands plants: the importance of dry matter content. AoB Plants, 8: pwl045. 10.1093/aobpla/plw045.

Bumb, I., Garnier, E., Coq, S., Nahmani, J., Del Rey Granado, M., Gimenez, O., Kazakou E (2018). Traits determining the digestibility– decomposability relationships in species from Mediterranean rangelands. Annals of Botany, 121(3): 459–469. 10.1093/aob/mcx175.

Chollet, S., Rambal, S., Fayolle, A., Hubert, D., Foulquié, D., & Garnier, E. (2014). Combined effects of climate, resource availability and plant traits on biomass produced in a Mediterranean rangeland. Ecology, 95: 737–748. 10.1890/13-0751.1.

*Cornwell, W.K., Cornelissen, J.H.C., Amatangelo, K., Dorrepaal, E., Eviner, V.T., Godoy, O., Hobbie, S.E., et al. (2008). Plant traits are the dominant control on litter decomposition rates within biomes worldwide. Ecology Letters, 11: 1065–1071. 10.1111/j.1461-0248.2008.01219.x.

Cortez, J., Garnier, E., Pérez-Harguindeguy, N., Debussche, M. & Gillon, D. (2007). Plant traits, litter quality and decomposition in a Mediterranean old-field succession. Plant and Soil, 296: 19–34. 10.1007/s11104-007-9285-6.

Delalandre, L., Violle, C., Coq, S., Garnier, E. (2023). Trait-environment relationships depend on species life history. Journal of Vegetation Science, 34(6): e13211. 10.1111/jvs.13211.

*Díaz, S., Kattge, J., Cornelissen, J.H.C., Wright, I.J., Lavorel, S., Dray, S., Reu, B., Kleyer, M., Wirth, C., Prentice, I.C., Garnier, E., et al. (2016). The global spectrum of plant form and function. Nature, 529: 167–171. 10.1038/nature16489.

*Díaz, S., Kattge, J., Cornelissen, J.H.C., Wright, I.J., Lavorel, S., Dray, S., Reu, B., Kleyer, M., Wirth, C., Prentice, I.C., Garnier, E., et al. (2022). The global spectrum of plant form and function: enhanced species-level trait dataset. Scientific Data, 9: 755. 10.1038/s41597-022-01774-9.

Dungan, R. J., Navas, M.-L., Duncan, R. P., et al. (2008). Effects of leaf emergence on leaf lifespan are independent of life form and successional status. Austral Ecology, 33: 932–939. 10.1111/j.1442-9993.2008.01875.x..

Fayolle, A., Violle, C., & Navas, M.-L. (2009). Differential impacts of plant interactions on herbaceous species recruitment: Disentangling factors controlling emergence, survival and growth of seedlings. Oecologia, 159: 817–825. https://doi.org/10.1007/s00442-008-1254-0

*Flores O, Garnier E, Wright IJ, Reich PB, Pierce S, Díaz S, Pakeman RJ, Rusch GM, Bernard-Verdier M, Testi B, et al. 2014. An evolutionary perspective on leaf economics: phylogenetics of leaf mass per area in vascular plants. Ecology and Evolution 4: 2799–2811. 10.1002/ece3.1087.

Fort F., Volaire F., Guilioni L., Barkaoui K., Navas M.-L., Roumet C. (2017). Root traits are related to plant water-use among rangeland Mediterranean species. Functional Ecology 31: 1700–1709. 10.1111/1365-2435.12888.

*Fortunel, C., Garnier, E., Joffre, R., Kazakou, E., Quested, H., Grigulis, K., Lavorel, S., et al. (2009). Leaf traits capture the effects of land use changes and climate on litter decomposability of grasslands across Europe. Ecology, 90: 598–611. 10.1890/08-0418.1

Fortunel, C., Violle, C., Roumet, C., Buatois, B., Navas, M.-L., Garnier, E. (2009). Allocation strategies and seed traits are hardly affected by nitrogen supply in 18 species differing in successional status. Perspectives in Plant Ecology, Evolution and Systematics, 11: 267–283. 10.1016/j.ppees.2009.04.003.

*Gardarin, A., Garnier, E., Carrère, P., et al. (2014). Plant trait–digestibility relationships across management and climate gradients in permanent grasslands. Journal of Applied Ecology, 51: 1207–1217. 10.1111/1365-2664.12293.

Garnier, E., Cortez, J., Billès, G., Navas, M.-L., Roumet, C., Debussche, M., Laurent, G., Blanchard, A., Aubry, D., Bellmann, A., Neill, C. & Toussaint, J.-P. (2004). Plant functional markers capture ecosystem properties during secondary succession. Ecology, 85: 2630–2637. 10.1890/03-0799.

Garnier, E., Fayolle, A., Navas, M.-L., Damgaard, C., Cruz, P., Hubert, D., Richarte, J., Autran, P., Leurent, C., Violle, C. (2018). Plant demographic and functional responses to management intensification: A long-term study in a Mediterranean rangeland. Journal of Ecology, 106: 1363–1376. 10.1111/1365-2745.12996.

Garnier, E., Laurent, G., Bellmann, A., Debain, S., Berthelier, P., Ducout, B., Roumet, C. & Navas, M.-L. (2001). Consistency of species ranking based on functional leaf traits. New Phytologist, 152: 69–83. 10.1046/j.0028-646x.2001.00239.x.

*Garnier, E., Lavorel, S., Ansquer, P., Castro, H., Cruz, P., Dolezal, J., Eriksson, O., et al. (2007). Assessing the effects of land use change on plant traits, communities and ecosystem functioning in grasslands: a standardized methodology and lessons from an application to 11 European sites. Annals of Botany, 99: 967–985. 10.1093/aob/mcl215.

Garnier, E., Vile, D., Debain, S., Bottin, L., Laurent, G. & Roumet, C. (2025). Photosynthesis, water-use and nitrogen relate to both plant height and leaf structure in 60 species from the Mediterranean. Functional Ecology 39: 567–582. doi: 10.1111/1365-2435.14737

Garnier, E., Vile, D., Roumet, C., Lavorel, S., Grigulis, K., Navas, M.-L., Lloret, F. (2019). Inter- and intra-specific trait shifts among sites differing in drought conditions at the north western edge of the Mediterranean Region. Flora, 254: 147–160. 10.1016/j.flora.2018.07.009.

Hummel I., Vile D., Violle C., Devaux J., Ricci B., Blanchard A., Garnier E., Roumet C. (2007). Relating root structure and anatomy to whole plant functioning in 14 herbaceous Mediterranean species. New Phytologist 173: 313–321. 10.1111/j.1469-8137.2006.01912.x.

*Kattge, J., Bönisch, G., Díaz, S., Lavorel, S., Prentice, I.C., Leadley, P., Tautenhahn, S., Werner, G.D.A., Aakala, T., Abedi, M., et al. (2020). TRY plant trait database – enhanced coverage and open access. Global Change Biology, 26: 119–188. 10.1111/gcb.14904.

*Kattge, J., Díaz, S., Lavorel, S., Prentice, I.C., Leadley, P., Bönisch, G., Garnier, E., et al. (2011). TRY – a global database of plant traits. Global Change Biology, 17: 2905–2935. 10.1111/j.1365-2486.2011.02451.x.

Kazakou, E, Vile, D., Shipley, B., Gallet, C. & Garnier, E. (2006). Co-variations in litter decomposition, leaf traits and plant growth in species from a Mediterranean old-field succession. Functional Ecology, 20: 21–30. 10.1111/j.1365-2435.2006.01080.x.

Kazakou, E., Bumb, I., Garnier, E. (2022). Species dominance rather than complementarity drives community digestibility and litter decomposition in species-rich Mediterranean rangelands. Applied Vegetation Science, 25(4): e12685. 10.1111/avsc.12685.

Kazakou, E., Garnier, E., Navas, M.-L., Roumet, C., Collin, C., Laurent, G. (2007). Components of nutrient residence time and the leaf economics spectrum in species from Mediterranean old-fields differing in successional status. Functional Ecology, 21: 235–245. 10.1111/j.1365-2435.2006.01242.x.

Kazakou, E., Gimenez, O., Garnier, E. (2007). Assessing the relative contribution of leaf lifespan and nutrient resorption to mean residence time: an elasticity analysis. Ecology, 88: 1857–1863. 10.1890/06-1352.1.

Kazakou, E., Violle, C., Roumet, C., Navas, M.-L., Vile, D., Kattge, J., Garnier, E. (2014). Are trait-based species rankings consistent across data sets and spatial scales? Journal of Vegetation Science, 25: 235–247. 10.1111/jvs.12066.

Kazakou, E., Violle, C., Roumet, C., Pintor, C., Gimenez, O., Garnier, E. (2009). Litter quality and decomposability of species from a Mediterranean succession depend on leaf traits but not on nitrogen supply. Annals of Botany, 104: 1151–116. 10.1093/aob/mcp202.

*Lavorel, S., de Bello, F., Grigulis, K., Lepš, J., Garnier, E., Castro, H., Dolezal, J., Godolets, C., Quétier, F. & Thébault, A. (2011). Response of herbaceous vegetation functional diversity to land use change across five sites in Europe and Israel. Israel Journal of Ecology and Evolution, 57: 53–72. 10.1560/IJEE.57.1-2.53.

Loranger, J., Blonder, B., Garnier, E., Shipley, B., Vile, D., Violle, C. (2016). Occupancy and overlap in trait space along a successional gradient in Mediterranean old fields. American Journal of Botany, 103: 1050–1060. 10.3732/ajb.1500483.

*Loranger, J., Violle, C., Shipley, B., Lavorel, S., Bonis, A., Cruz, P., Louault, F., Loucougaray, G., Mesléard, F., Yavercovski, N., et al. (2016). Recasting the dynamic equilibrium model through a functional lens: the interplay of trait-based community assembly and climate. Journal of Ecology, 104: 781–791. 10.1111/1365-2745.12536.

Navas, M.-L. Ducout, B., Roumet, C., et al. (2003). Leaf life span, dynamics and construction cost of species from Mediterranean old-fields differing in successional status. New Phytologist, 159 (1): 213–228. 10.1046/j.1469-8137.2003.00790.x.

Navas, M.-L., Roumet, C., Bellmann, A., et al. (2010). Suites of plant traits in species from different stages of a Mediterranean secondary succession. Plant Biology, 12: 183–196. 10.1111/j.1438-8677.2009.00208.x.

*Pakeman, R.J., Garnier, E., Lavorel, S., Ansquer, P., Castro, H., Cruz, P., Doležal, J., et al. (2008). Impact of abundance weighing on the response of seed traits to climate and land use. Journal of Ecology, 96: 355–366. 10.1111/j.1365-2745.2007.01336.x.

*Pakeman, R.J., Lepš, J., Kleyer, M., Lavorel, S., Garnier, E., VISTA Consortium. (2009). Relative climatic, edaphic and management controls of plant functional trait signatures. Journal of Vegetation Science, 20: 148–159. 10.1111/j.1654-1103.2009.05548.x

Pérez-Ramos, I.M., Roumet, C., Cruz, P., Blanchard, A., Autran, P. & Garnier, E. (2012). Evidence for a « plant community economics spectrum » driven by nutrient and water limitations in a Mediterranean rangeland of Southern France. Journal of Ecology, 100: 1315–1327. 10.1111/1365-2745.12000.

Poirier V., Roumet C., Angers D.A., Munson A.D. (2018). Species and root traits impact macroaggregation in the rhizospheric soil of a Mediterranean common garden experiment. Plant and Soil, 424: 289–302. 10.1007/s11104-017-3407-6.

Prieto I, Birouste M., Zamora-Ledezma E., Gentit A., Goldin J., Volaire F., Roumet C. (2017). Decomposition rates of fine roots from three herbaceous perennial species: combined effect of root mixture composition and living plant community. Plant and Soil, 415: 359–372. 10.1007/s11104-016-3163-z.

Prieto I., Querejeta J., Segrestin J., Volaire F., Roumet C. (2018). Leaf carbon and oxygen isotopes are coordinated with the leaf economics spectrum in Mediterranean rangeland species. Functional Ecology, 32: 612–625. 10.1111/1365-2435.13025.

Roumet C., Birouste M., Picon-Cochard C., Ghestem M., Osman N., Vrignon-Brenas S., Cao K., Stokes A. (2016). Root structure - function relationships in 74 herbaceous species: evidence of a root economics spectrum related to carbon economy. New Phytologist, 210: 815–826. 10.1111/nph.13828.

Segrestin, J., Bernard-Verdier, M., Violle, C., Richarte, J., Navas, M.-L., & Garnier, E. (2018). When is the best time to flower and disperse? A comparative analysis of plant reproductive phenology in the Mediterranean. Functional Ecology, 32: 1770–1783. 10.1111/1365-2435.13098.

Segrestin, J., Kazakou, E., Coq, S., Sartori, K., Richarte, J., Rowe, N.P. & Garnier, E. (2023). Responses of leaf biomechanics and underlying traits to rangeland management differ between graminoids and forbs. Journal of Vegetation Science, 34(6): e13216. 10.1111/jvs.13216.

Segrestin, J., Navas, M.-L., & Garnier, E. (2020). Reproductive phenology as a dimension of the phenotypic space in 139 plant species from the Mediterranean. New Phytologist, 225: 740–753. 10.1111/nph.16165.

Segrestin, J., Sartori, K., Navas, M.-L., Kattge, J., Díaz, S., Garnier, E. (2021). PhenoSpace: A Shiny application to visualize trait data in the phenotypic space of the global spectrum of plant form and function. Ecology and Evolution, ece3.6928. 10.1002/ece3.6928.

Shipley, B., Vile, D. & Garnier, E. (2006). From plant traits to plant communities: a statistical mechanistic approach to biodiversity. Science, 314: 812–814. 10.1126/science.1131344.

*Vile, D., Garnier, E., Shipley, B., Laurent, G., Navas, M.-L., Roumet, C., Lavorel, S., Díaz, S., Hodgson, J.G., Lloret, F., Midgley, G.F., Poorter, H., Rutherford, M.C., Wilson, P.J. & Wright, I.J. (2005). Specific leaf area and dry matter content estimate thickness in laminar leaves. Annals of Botany, 96: 1129–1136. 10.1093/aob/mci264.

Vile, D., Shipley, B. & Garnier, E. (2006). A structural equation model to integrate changes in functional strategies during old-field succession. Ecology, 87: 504–517. 10.1890/05-0822.

Vile, D., Shipley, B. & Garnier, E. (2006). Ecosystem productivity can be predicted from potential relative growth rate and species abundance. Ecology Letters, 9: 1061–1067. 10.1111/j.1461-0248.2006.00958.x.

Violle, C., Castro, H., Richarte, J., & Navas, M. (2009). Intraspecific seed trait variations and competition: Passive or adaptive response? Functional Ecology, 23: 612–620. 10.1111/j.1365-2435.2009.01539.x

Violle C., Garnier E., Lecoeur J., Roumet C., Podeur C., Blanchard A., Navas M.-L. (2009). Competition, traits and resource depletion in plant communities. Oecologia, 160:747–755. 10.1007/s00442-009-1333-x.

*Wright, I.J., Reich P.B., Cornelissen, J.H.C., Falster, D.S., Garnier, E., Hikosaka, K., Lamont, B.B., Lee, W., Oleksyn, J., Osada, N., Poorter, H., Villar, R., Warton, D.I., Westoby, M. (2005). Assessing the generality of global leaf trait relationships. New Phytologist, 166: 485–496. 10.1111/j.1469-8137.2005.01349.x.

*Wright, I.J., Reich, P.B., Westoby, M., Ackerly, D.D., Baruch, Z., Bongers, F., Cavender-Bares, J., et al. (2004). The worldwide leaf economics spectrum. Nature, 428: 821–827. 10.1038/nature02403.

## Literature Citations

Anderegg, L. D. L. 2023. Why can’t we predict traits from the environment? New Phytologist 237:1998–2004.

Barkaoui, K., C. Roumet, and F. Volaire. 2016. Mean root trait more than root trait diversity determines drought resilience in native and cultivated Mediterranean grass mixtures. Agriculture, Ecosystems & Environment 231:122–132.

Bernard, C. 2008. Flore des Causses. Hautes terres, gorges, vallées et vallons (Aveyron, Lozère, Hérault et Gard). Bulletin de la Société Botanique du Centre-Ouest (Nouvelle série) 31:1–784.

de Bolòs, O., J. Vigo, R. Masalles, and J. Ninot. 1993. Flora manual dels països catalans. Second. Editorial Pòrtic s.a., Barcelona.

Braun-Blanquet, J., N. Roussine, and R. Nègre. 1952. Les Groupements Végétaux de la France Méditerranéenne. Centre National de la Recherche Scientifique, Montpellier.

Cornelissen, J. H. C., S. Lavorel, E. Garnier, S. Díaz, N. Buchmann, D. E. Gurvich, P. B. Reich, H. ter Steege, H. D. Morgan, M. G. A. van der Heijden, J. G. Pausas, and H. Poorter. 2003. A handbook of protocols for standardised and easy measurement of plant functional traits worldwide. Australian Journal of Botany 51:335–380.

Coste, A. H. 1937. Flore descriptive et illustrée de la France, de la Corse et des contrées limitrophes. Librairie des Sciences et des Arts, Paris, France.

Daget, P. 1977. Le bioclimat Méditerranéen: analyse des formes climatiques par le système d’Emberger. Vegetatio 34:87–103.

Ely, K. S., A. Rogers, D. A. Agarwal, E. A. Ainsworth, L. P. Albert, A. Ali, J. Anderson, M. J. Aspinwall, C. Bellasio, C. Bernacchi, S. Bonnage, T. N. Buckley, J. Bunce, A. C. Burnett, F. A. Busch, A. Cavanagh, L. A. Cernusak, R. Crystal-Ornelas, J. Damerow, K. J. Davidson, M. G. De Kauwe, M. C. Dietze, T. F. Domingues, M. E. Dusenge, D. S. Ellsworth, J. R. Evans, P. P. G. Gauthier, B. O. Gimenez, E. P. Gordon, C. M. Gough, A. H. Halbritter, D. T. Hanson, M. Heskel, J. A. Hogan, J. R. Hupp, K. Jardine, J. Kattge, T. Keenan, J. Kromdijk, D. P. Kumarathunge, J. Lamour, A. D. B. Leakey, D. S. LeBauer, Q. Li, M. R. Lundgren, N. McDowell, K. Meacham-Hensold, B. E. Medlyn, D. J. P. Moore, R. Negrón-Juárez, Ü. Niinemets, C. P. Osborne, A. L. Pivovaroff, H. Poorter, S. C. Reed, Y. Ryu, A. Sanz-Saez, S. C. Schmiege, S. P. Serbin, T. D. Sharkey, M. Slot, N. G. Smith, B. V. Sonawane, P. F. South, D. C. Souza, J. R. Stinziano, E. Stuart-Haëntjens, S. H. Taylor, M. D. Tejera, J. Uddling, V. Vandvik, C. Varadharajan, A. P. Walker, B. J. Walker, J. M. Warren, D. A. Way, B. T. Wolfe, J. Wu, S. D. Wullschleger, C. Xu, Z. Yan, and D. Yang. 2021. A reporting format for leaf-level gas exchange data and metadata. Ecological Informatics 61:101232.

Escarré, J., C. Houssard, and M. Debussche. 1983. Evolution de la végétation et du sol après abandon cultural en région méditerranéenne : étude de succession dans les garrigues du Montpelliérais (France). Acta Oecologica, Oecologia Plantarum 4:221–239.

Fort, F., F. Volaire, L. Guilioni, K. Barkaoui, M.-L. Navas, and C. Roumet. 2017. Root traits are related to plant water-use among rangeland Mediterranean species. Functional Ecology 31:1700–1709.

Freschet, G. T., L. Pagès, C. M. Iversen, L. H. Comas, B. Rewald, C. Roumet, J. Klimešová, M. Zadworny, H. Poorter, J. A. Postma, T. S. Adams, A. Bagniewska-Zadworna, A. G. Bengough, E. B. Blancaflor, I. Brunner, J. H. C. Cornelissen, E. Garnier, A. Gessler, S. E. Hobbie, I. C. Meier, L. Mommer, C. Picon-Cochard, L. Rose, P. Ryser, M. Scherer-Lorenzen, N. A. Soudzilovskaia, A. Stokes, T. Sun, O. Valverde-Barrantes, M. Weemstra, A. Weigelt, N. Wurzburger, L. M. York, S. Batterman A., M. Gomes de Moraes, Š. Janeček, H. Lambers, V. Salmon, N. Tharayil, and M. L. McCormack. 2021. A starting guide to root ecology: strengthening ecological concepts and standardising root classification, sampling, processing and trait measurements. New Phytologist 232:973– 1122.

Garnier, E., J. Cortez, G. Billès, M.-L. Navas, C. Roumet, M. Debussche, G. Laurent, A. Blanchard, D. Aubry, A. Bellmann, C. Neill, and J.-P. Toussaint. 2004. Plant functional markers capture ecosystem properties during secondary succession. Ecology 85:2630– 2637.

Garnier, E., G. Laurent, A. Bellmann, S. Debain, P. Berthelier, B. Ducout, C. Roumet, and M.-L. Navas. 2001. Consistency of species ranking based on functional leaf traits. New Phytologist 152:69–83.

Garnier, E., S. Lavorel, P. Ansquer, H. Castro, P. Cruz, J. Dolezal, O. Eriksson, C. Fortunel, H. Freitas, C. Golodets, K. Grigulis, C. Jouany, E. Kazakou, J. Kigel, M. Kleyer, V. Lehsten, J. Lepš, T. Meier, R. J. Pakeman, M. Papadimitriou, V. . P. Papanastasis, H. Quested, F. Quétier, M. Robson, C. Roumet, G. Rusch, M. Skarpe, M. Sternberg, J.-P. Theau, A. Thébault, D. Vile, and M. Zarovali. 2007. Assessing the effects of land use change on plant traits, communities and ecosystem functioning in grasslands: a standardized methodology and lessons from an application to 11 European sites. Annals of Botany 99:967–985.

Garnier, E., M.-L. Navas, and K. Grigulis. 2016. Plant Functional Diversity - Organism Traits, Community Structure, and Ecosystem Properties. Oxford University Press, Oxford.

Garnier, E., U. Stahl, M.-A. Laporte, J. Kattge, I. Mougenot, I. Kühn, B. Laporte, B. Amiaud, F. S. Ahrestani, G. Bönisch, D. E. Bunker, J. H. C. Cornelissen, S. Díaz, B. J. Enquist, S. Gachet, P. Jaureguiberry, M. Kleyer, S. Lavorel, L. Maicher, N. Pérez-Harguindeguy, H. Poorter, M. Schildhauer, B. Shipley, C. Violle, E. Weiher, C. Wirth, I. J. Wright, and S. Klotz. 2017. Towards a thesaurus of plant characteristics: an ecological contribution. Journal of Ecology 105:298–309.

Kazakou, E., E. Garnier, M.-L. Navas, C. Roumet, C. Collin, and G. Laurent. 2007. Components of nutrient residence time and the leaf economics spectrum in species from Mediterranean old-fields differing in successional status. Functional Ecology 21:235–245.

Keller, A., M. J. Ankenbrand, H. Bruelheide, S. Dekeyzer, B. J. Enquist, M. B. Erfanian, D. S. Falster, R. V. Gallagher, J. Hammock, J. Kattge, S. D. Leonhardt, J. S. Madin, B. Maitner, M. Neyret, R. E. Onstein, W. D. Pearse, J. H. Poelen, R. Salguero-Gomez, F. D. Schneider, A. B. Tóth, and C. Penone. 2023. Ten (mostly) simple rules to future-proof trait data in ecological and evolutionary sciences. Methods in Ecology and Evolution 14:444–458.

Le Floc’h, E., J. Aronson, S. Dhillion, J.-L. Guillerm, A. Grossmann, and E. Cunge. 1998. Biodiversity and ecosystem trajectories: first results from a new LTER in southern France. Acta Oecologica 19:285–293.

Lloret, F., and M. Vilá. 2003. Diversity patterns of plant functional types in relation to fire regime and previous land use in Mediterranean woodlands. Journal of Vegetation Science 14:387–398.

McCormack, M. L., D. Guo, C. M. Iversen, W. Chen, D. M. Eissenstat, C. W. Fernandez, L. Li, C. Ma, Z. Ma, H. Poorter, et al. 2017. Building a better foundation: improving root-trait measurements to understand and model plant and ecosystem processes. New Phytologist 215:27–37.

Molénat, G., D. Foulquié, P. Autran, J. Bouix, D. Hubert, M. Jacquin, F. Bocquier, and B. Bibé. 2005. Pour un élevage ovin allaitant performant et durable sur parcours: un système expérimental sur le Causse du Larzac. INRA Productions Animales 18:323–338.

Mungall, C. J., G. V. Gkoutos, C. L. Smith, M. A. Haendel, S. E. Lewis, and M. Ashburner. 2010. Integrating phenotype ontologies across multiple species. Genome Biology 11:R2.

Navas, M.-L., C. Roumet, A. Bellmann, G. Laurent, and E. Garnier. 2010. Suites of plant traits in species from different stages of a Mediterranean secondary succession. Plant Biology 12:183–196.

Pakeman, R. J., and Quested, H. M. 2007. Sampling plant functional traits: What proportion of the species need to be measured? Applied Vegetation Science 10: 91–96.

R Core Team 2020.

Raunkiaer, C. 1934. The Life Forms of Plants and Statistical Plant Geography. Page (H. Gilbert-Carter, Tran.). English Edition. Oxford University Press, Oxford.

Schneider, F. D., D. Fichtmueller, M. M. Gossner, A. Güntsch, M. Jochum, B. König-Ries, G. Le Provost, P. Manning, A. Ostrowski, C. Penone, and N. K. Simons. 2019. Towards an ecological trait-data standard. Methods in Ecology and Evolution 10:2006–2019.

Shipley, B., F. De Bello, J. H. C. Cornelissen, E. Laliberté, D. C. Laughlin, and P. B. Reich. 2016. Reinforcing loose foundation stones in trait-based plant ecology. Oecologia 180:923–931.

Tavşanoğlu, Ç., and J. G. Pausas. 2018. A functional trait database for Mediterranean Basin plants. Scientific Data 5:180135.

Tison, J.-M., P. Jauzein, H. Michaud, D. Jeanmono, and F. Boillot. 2014. Flore de la France méditerranéenne continentale. 1ère édition. Naturalia Publications, Turriers Porquerolles.

Violle, C., M.-L. Navas, D. Vile, E. Kazakou, C. Fortunel, I. Hummel, and E. Garnier. 2007. Let the concept of trait be functional! Oikos 116:882–892.

Walter, H., E. Harnickell, and D. Mueller-Dombois. 1975. Climate-diagram maps of the individual continents and the ecological climatic regions of the Earth. Springer-Verlag, Berlin.

Wilkinson, M. D., M. Dumontier, Ij. J. Aalbersberg, G. Appleton, M. Axton, A. Baak, N. Blomberg, J.-W. Boiten, L. B. da Silva Santos, P. E. Bourne, J. Bouwman, A. J. Brookes, T. Clark, M. Crosas, I. Dillo, O. Dumon, S. Edmunds, C. T. Evelo, R. Finkers, A. Gonzalez-Beltran, A. J. G. Gray, P. Groth, C. Goble, J. S. Grethe, J. Heringa, P. A. C. ’t Hoen, R. Hooft, T. Kuhn, R. Kok, J. Kok, S. J. Lusher, M. E. Martone, A. Mons, A. L. Packer, B. Persson, P. Rocca-Serra, M. Roos, R. van Schaik, S.-A. Sansone, E. Schultes, T. Sengstag, T. Slater, G. Strawn, M. A. Swertz, M. Thompson, J. van der Lei, E. van Mulligen, J. Velterop, A. Waagmeester, P. Wittenburg, K. Wolstencroft, J. Zhao, and B. Mons. 2016. The FAIR Guiding Principles for scientific data management and stewardship. Scientific Data 3:160018.

WRB. 2022. World Reference Base for Soil Resources. IUSS Working Group WRB 2022. ISBN 979-8-9862451-1-9

